# Single-molecule analysis of synaptic protein complexes and vesicle recruitment

**DOI:** 10.1101/2025.08.19.671146

**Authors:** Akshay Kapadia, Anne-Sophie Hafner

**Author notes:** Technical contact (Akshay Kapadia).

## Abstract

Single-molecule pull-down (SIM-Pull) combined with TIRF microscopy enables direct visualization of proteins and multi-protein complexes. Here, we present an extended SIM-Pull protocol for analyzing protein interactions at the active zone and their ability to recruit isolated synaptic vesicles (SV). SV recruitment mediated by STX1A-SNARE or RIM1-Rab3a interactions, respectively; can be directly visualized and quantified. This technique opens new avenues to examine the subcellular vesicle-associated protein-protein interactions at a molecular level in a near-native cellular context.

**Highlights:**

- Extended SIM-Pull protocol combining biochemical isolation with TIRF microscopy to study synaptic protein complexes at near-native environment
- Enables direct quantification of synaptic vesicle recruitment at the surface (active zone) *via* protein-mediated vesicle tethering
- Adaptable platform for probing molecular interactions of protein complexes within different neuronal, cellular or subcellular compartments

**Graphical abstract:** 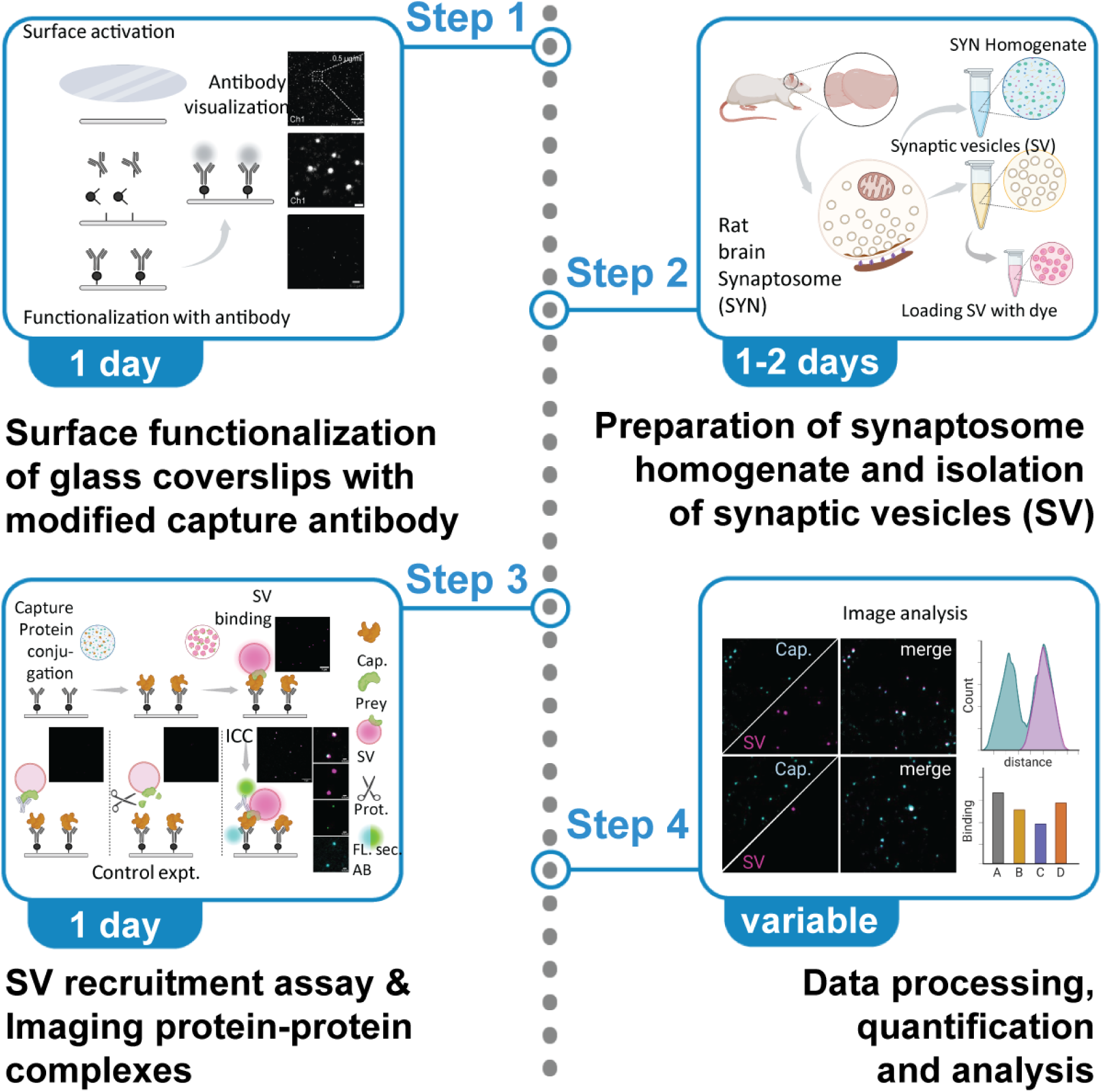

**Institutional permissions:** Animals were handled and maintained according to the guidelines laid down by the Animal Welfare Body (AWB) (Instantie voor Dierenwelzijn IvD) in line with the animal experimentation policy within Radboud University and RadboudUMC; under the license/protocol numbers 2021-0040-001/002 to Dr. Anne-Sophie Hafner.

## Innovation

SIM-Pull technique combines immunoisolation with single-molecule TIRF imaging to directly visualize and quantify protein–protein interactions at nanometer resolution. This powerful approach enables detection of complex stoichiometry, binding dynamics, and functional assembly in near-native molecular environments (1–7).

- Here, we describe a streamlined workflow protocol to visualize synaptic vesicle (SV) recruitment through defined protein–protein interactions in a near-native environment. By combining antibody-based immobilization of capture proteins (STX1A or RIM1), the application of fluorescently labeled native SVs containing prey proteins (SNAREs or Rab3a), allowing real-time visualization of vesicle tethering events *via* TIRF microscopy.
- The method enables quantitative assessment of binding specificity, vesicle recruitment efficiency, and the effects of epitope blocking or proteolytic disruption. Unlike prior approaches relying on reconstituted vesicles or detergent-extracted components, this assay preserves native vesicle composition and topology, providing biologically relevant insights into protein-protein interactions.
- The modular format also supports parallel testing of multiple interacting partners and mutant variants, making it a powerful platform for probing synaptic protein function and vesicle targeting under physiologically relevant conditions.
- This technology lays the groundwork for dissecting vesicle recruitment mechanisms across diverse cellular systems and could be readily adapted to study subcellular vesicle docking or trafficking events by facilitating protein interactions in a controlled, quantifiable, and near-native environment.

## Before you begin

- TIRF microscope and imaging setup

- Ensure that the microscope and laser lines are installed on an air-gapped table to limit vibration that will impact image acquisition.
- Always check laser alignment to achieve TIRF mode prior to imaging sessions.
- The microscope is equipped with a 100-x objective (N/A 1.4 or higher) and a DS-Fi3 (Nikon) video camera (or equivalent) for fast image acquisition.
- The protocol described herewith uses 25mm glass coverslips, requiring the use of quick release magnetic and non-magnetic imaging chambers suitable for a 25 mm coverslip - imaging aperture of 19.7 mm, imaging buffer volume of 308 µl (Warner Instruments, #QR-40LP). This was staged within a culture dish microincubator (Warner instruments, #DH-40iL) **Note:** If you would like to procure a new setup, we recommend the open round bath perfusion integrated imaging chamber (Warner instruments, #RC-21BRW), suitable for 25 mm glass coverslips.
- The perfusion system on the microincubator was connected to a syringe pump (Fusion 100-X Touch infusion syringe pump, KR Analytics) to facilitate flow of buffers/solutions during washing and incubation steps. **Note:** This is optional, adding and removing reagents/buffers could also be done carefully by using a pipette.
- Another alternative is the use of Ibidi μ-dish Grid-500 glass bottom (Ibidi # 81168). The pre-etched grid on the glass surface will ease quantification.

## Antibody thiolation and surface functionalization

This protocol enables the covalent immobilization of antibodies onto glass surfaces, allowing for stable, specific, and reproducible capture of target molecules for molecular interaction studies (8,9). This method results in a low-background, high-specificity functionalized surface ideal for single-molecule imaging and detection applications. This optimized SATA-based thiolation approach (10,11) was found to be superior to partial reduction strategies [e.g., DTT or 2-MEA (12,13)], which caused loss of antibody activity.

You can also follow other surface activation protocols previously described in literature.

**Timing: [1 day, variable]**

1. Addition of protected Sulfhydryl groups to the antibody (Fig. 1A, *scheme*)

a. Dilute or dissolve 10-20 μg antibody in 200 μl of phosphate buffer. **Note:** We recommend doing a pilot run through with a small volume of the antibody you would like to test with this protocol. Please ensure that timing of reactions, reactivity of antibodies post modifications are comparable or within detectable range to the non-modified antibody. Starting concentration of the antibody should be calculated based on the desired final concentration for each coverslip, multiplied to the number of coverslips to include all testing conditions + 4(factors in the loss during purification and desalting steps + samples for analysis). Additionally, this protocol has been optimized for IgG antibodies, but could be extended to IgA and IgMs, following pilot optimization steps.
b. Prepare 15-20 μM solution of SATA in DMSO. Prepare only as much as needed as the rest of the unused solution will be discarded. Immediately add 20 μl SATA solution to the antibody solution (ensure > 20-fold molar excess of reagent).
c. Incubate for 30-60 min at room temperature (RT, 20-25°C).
d. Purify the modified antibody-**S–Acetyl** from excess reaction by-products using a desalting column equilibrated with phosphate buffer (pH 7.2). Collect 50 μl fractions in fresh tubes. **Note:** We prepared a desalting column using a low-bind 2.5 mL dispensing quick-tip to purify/collect smaller volumes of antibody solutions.
e. Identify the fraction(s) containing antibody (absorbance at λ 280 nm) and pool the fractions with the antibody (optional WB + ELISA analysis). **Note:** <u>Stopping point</u>, the modified antibody may be stored indefinitely at -20°C, as reported in literature, but we always proceeded to step 3 immediately, or the next day.
2. Deprotection with Hydroxylamine (Fig. 1A, *scheme*) **Critical:** Before starting with this step, make sure you have already completed step 3.
  a. Dissolve 3.48 mg of Hydroxylamine hydrochloride in 0.8 ml phosphate buffer and then adjust to pH 7.2 with NaOH. Finally adjust the volume to 1 ml with phosphate buffer (final concentration ∼0.5 M Hydroxylamine hydrochloride). **Note:** Since these are small volumes, we recommend using pH strips.
  b. Add Hydroxylamine solution to the SATA-modified antibody solution in the ratio of 1:9.
  c. Incubate for 2 h at RT, mild agitation. (Thermomix, without temperature control, 200 rpm)
  d. Purify the antibody from the excess hydroxylamine using a desalting Column equilibrated with phosphate buffer (pH 7.2). Follow steps as described in 1d-e. Collect 50 μl fractions. **Critical:** Once the steps for deprotection (i.e. deacetylation) of the sulfhydryl groups are performed, the following steps need to be performed as quickly as possible. The antibody must be desalted again and used immediately for coupling to the surface-modified glass coverslips.
  e. Identify the fraction(s) that contain antibodies by measuring those having peak absorbance at λ 280 nm. Pool the fractions that contain antibodies. **Important:** You need to work swiftly and proceed immediately to section 4. **Note:** To ensure that your antibody is still functional we recommend running a western blot -WB and/or ELISA panel, *e.g.* Coomassie staining to check for the migratory patterns of the heavy and light chains of the modified antibody in denaturation conditions (Fig. 1Bi and Fig. S1i), b. western blotting (Fig. 1Bi-ii and Fig. S1i-ii) or using modified ELISA (Fig. 1Bii and Fig. S1Aii). We also recommend using an indirect ELISA panel to re-examine the LOD of the antibody detecting properties (Fig. 1C and Fig. S1B). In our experiments we observed differences in the LOD levels pointing towards the fact that the modification of the antibody slightly alters the epitope recognition site (Fig. 1C and Fig. S1B, differently affect different antibody subtypes). For indirect and sandwich ELISA, please refer to literature published methods (14) or protocols followed in this study (detailed description in SI, section 1A).
3. Surface activation of glass coverslips (Fig. S2A, *scheme*)
  a. Clean coverslips thoroughly in 100% ethanol and air dry over a particle/dust-free tissue. These coverslips can be stored in sealed, cool and desiccated conditions for indefinite time. **Critical:** You should handle coverslips carefully with powder-free gloves and try to work in a clean and particle/dust free environment as much as possible for the entirety of this step. **Note:** Literature reports recommend baking the coverslips and/or cleaning surfaces with plasma or UV-ozone treatment prior to functionalization; but we did not observe key specific differences for the antibodies tested in our system, so we continued with coverslips that were only ethanol-treated and air-dried.
  b. In a fume hood, prepare 2% APTES (3-Aminopropyltriethoxysilane) in dry acetone in a wide base and tall glass beaker. **Critical:** This one-time use solution should always be prepared fresh prior to coverslip treatment. Please ensure acetone is free of water; you can use desiccant beads or purge the acetone with inert N_2_ gas for 30 min prior to use. If you have decided to use 35-mm glass bottom dishes, please test its compatibility with acetone before starting the experiment. **Caution!** APTES and Acetone handling requires special precautions, please refer to the materials section and ensure all safety precautions are following during this step.
  c. Next, immerse coverslips for 45 s - 1 min, rinse (3 x) in dry acetone, and air-dry over a particle/dust-free tissue. **Note:** Coverslips can be stored together in a petri dish or individually in a 6 well multi-plate. These coverslips can be stored indefinitely under desiccated conditions.
  d. Prepare phosphate buffer (1x PBS in ultrapure water - Milli-Q, pH 7.2 supplemented with 10 mM EDTA). Filter through 0.22 μm microfilter. **Note:** You can prepare 1L of 5x solution, aliquot into 50 mL portions and store at 4C for no longer than 1 year. **Critical:** Please always use filtered ultrapure water to prepare all the buffers in this experimental protocol. It is optional to autoclave water before use.
  e. Prepare Sulfo-SMCC (0.5-2 mg/mL) fresh in phosphate buffer (adjust pH to 7.5, important). Always prepare fresh and use the entire solution, do not store the solution for repeated/ later use.
  f. Add the solution to coverslips (ensure that the entire coverslip is dipped into the solution and there are no dry spots) and incubate with agitation for 2 h at RT with rotational agitation. **Note:** We observed the speed of agitation should be optimal (not be very slow and not very fast). Orbital rotational speed between 40-60 rpm and linear agitation speed between 100-120 rpm. We recommend using a circular rotational shaker resulting in uniform coating and distribution of antibody molecules.
  g. Rinse with phosphate buffer thoroughly (2 x quick washes + 2 x washes with 5 min interval). Remove the solution as much as possible, to proceed to functionalization with the modified antibody (step 4). (<u>Stopping point</u>) Plates can be stored for a year at 4°C in dark desiccated conditions. **Note:** Until this step, coverslips can be prepared in bulk and stored until use. Before starting with step 4, please ensure enough coverslips to include all testing conditions + an additional 2 *(factors in the loss of coverslips during handling).
4. Surface functionalization *via* cross-linking of reactive Sulfhydryl-antibody (Figure 1A and 2A, *schemes*)
  a. Preincubate the glass coverslips with phosphate buffer for 10 min, discard the solution and repeat this step one additional time. **Note:** Begin this step just before starting step 2d.
  b. Cover the maleimide-activated surface material with the antibody solution (prepared above in step 2e and further diluted in phosphate buffer to a volume sufficient to cover the entire surface.) **Critical:** For optimal results, ensure that the final antibody concentration is within the range of 0.5-1 μg/ml (Figure S2B-D).
  c. Incubate for 2-4 h at RT with optimal agitation. Please see section 3f. **Note:** The time must be determined based on the sensitivity of the antibody. A clear idea could be derived from the LOD values of the specific antibody. In our hands, lower LOD values, lower incubation duration.
  d. Remove the antibody solution and thoroughly rinse the surface with phosphate buffer (2 x quick washes + 2 x washes with 5 min interval) to ensure that only covalently attached antibody molecules remain.
  e. The surface is now ready to use for detection assays and other applications. You can store this antibody coated glass coverslips in phosphate buffer containing 0.01-0.02% sodium azide at 4°C for 1-2 months (storage time could be extended until 6 months). **Note:** Successful completion of this step will yield well-resolved and well-distributed antibody molecules (Fig. 3A). Since we wanted to test the recruitment of synaptic vesicles we decided to use Syntaxin-1 (STX1) - SNAREs (Synaptobrevins-Syb) and Rab3-interacting molecule 1 (RIM1) - Ras-related protein Rab-3A, interactions which are well documented in literature (15–17). Using this protocol have successfully used the following antibodies: Anti-STX1A (Clone: 230470F5, rabbit polyclonal, IgG; ProteinTech #83159-6-RR) and Anti-RIM1/2 (Clone: SY-53E12, rat monoclonal, IgG2a-κ light chain; Synaptic Systems #140217).
5. Testing the quality of functionalized coverslips
  a. Assemble the coverslip in the imaging chamber and apply 400-500 μl of imaging buffer and acquire 10-15 images at suitable laser power image at the TIRF microscope.
    **Critical:** It is recommended to record the background for at least 1 coverslip from each batch of surface functionalization of coverslips (Please see Fig. S2, C-D).
    **Note:** You can store this coverslip and image the background fluorescence during major step 2-2e. Please note, for quantification it is important to use images obtained from the antigen delete control conditions in major steps 1-3a and 1b.
  b. The average number of fluorescent molecules per unit imaging area, is your background fluorescence. Ideally, the observed background fluorescence is <0.02 molecules per μm^2^ (or < 5 particles per FOV under our experimental conditions). Representative images have been depicted in Fig. 1D.
6. Alternate protocols: Antibody Thiolation *via* Partial Reduction with 2-MEA or DTT Alternatively, to generate free thiol groups for antibody immobilization, we tested two conventional partial reduction methods.
  a. Antibodies were diluted in phosphate buffer and treated with freshly prepared 0.25 M β-ME (β-mercaptoethanol) or 0.5 M DTT (dithiothreitol) at 37°C for 90 min (Fig. 1b and S1A).
  b. Following incubation step 6a, samples were desalted, and peak fractions (absorbance at λ 280) were pooled as indicated earlier. **Note:** Neither methods yielded satisfactory results in our hands as we observed loss of antibody activity after 90 min of incubation (Fig. 1Bi-ii and Fig. S1Ai-ii) as well as short time points (Fig. S1C). Possible issues could be due to incomplete reduction or excessive modification of the parent antibody. We therefore discontinued their use in favor of SATA-based thiolation methods described above, which provided more consistent and functionally viable antibody conjugates in our system.

**Figure 1.**
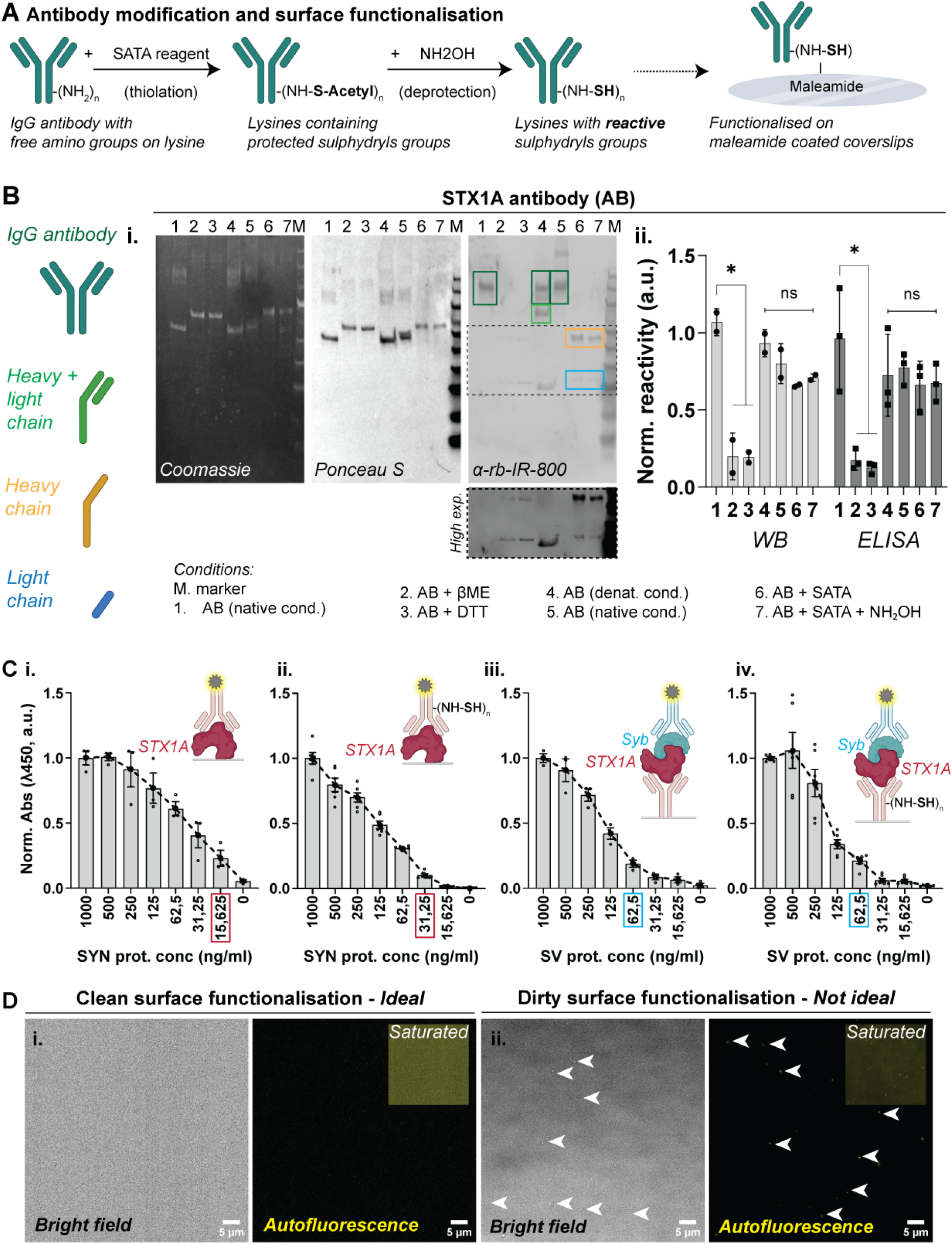
Antibody thiolation and validation of capture efficiency. (A) Schematic of antibody thiolation and functionalization workflow: IgG antibodies are modified using SATA to introduce protected sulfhydryl groups, which are subsequently deprotected with hydroxylamine (NH₂OH) to yield reactive thiols for conjugation to maleimide-coated surfaces. (B) Validation of antibody reactivity post-thiolation. (i) SDS-PAGE under various reducing and non-reducing conditions showing heavy and light chains detected by Coomassie, Ponceau, and western blot (WB) with anti-rabbit secondary. N = 2. (ii) Quantification of antibody reactivity by WB and modified ELISA reveals that SATA thiolation (condition 6) did not significantly impair antibody function, and reactivity is preserved after deprotection (condition 7). WB, N = 2, ELISA, N = 3. One-way ANOVA, * *p* < 0.05. (C) Syntaxin1 antibody reactivity in indirect and sandwich ELISA assays. (i–ii) STX1A antibody with or without thiolation detects STX1A protein in synaptosome lysates in a dose-dependent manner; examined using indirect ELISA. (iii) Sandwich ELISA showing capture of synaptobrevin (Syb) to STX1A protein immobilized *via* antibody without (iii; N = 2, n = 4) and with thiolation steps (iv; N = 3, n = 6). (D) Representative bright field and autofluorescence (488 laser channel, *yellow*) images showing ideal surface activated coverslips with negligible background. Improper surface activation, using non-filtered buffer or unclean conditions result in excessive background which could alter results.

**Figure 2.**
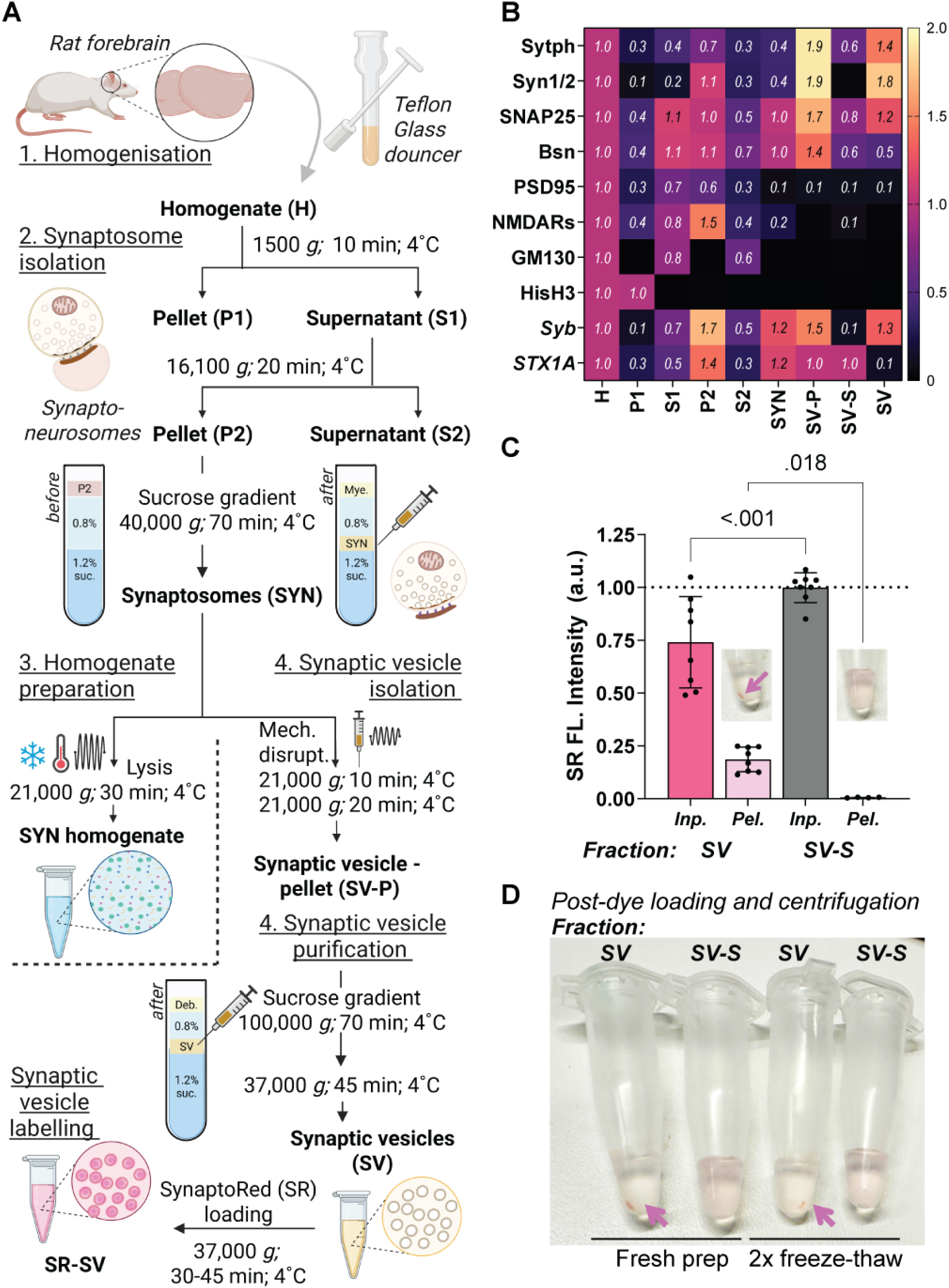
Workflow for preparation of synaptosome homogenate, synaptic vesicle isolation and purification from rat forebrain. (A) Schematic overview of the workflow using Syn-PER buffer. Rat forebrains are homogenized using a glass-Teflon douncer, followed by differential centrifugation to isolate synaptosomes (SYN) *via* a discontinuous sucrose gradient (Step 1–2). SYN fraction is lysed (Step 3) or disrupted to release vesicles (Step 4). Synaptic vesicles are further purified *via* centrifugation and a sucrose gradient step and purified SVs labelled with SynaptoRed (SR) for visualization. (B) Heatmap showing relative enrichment (normalized values) of synapse and synaptic vesicle specific proteins across fractions from the workflow (H, P1, S1, P2, S2, SYN, SV-P, SV-S, and SV). Values are normalized to homogenate. SV fractions are highly enriched in presynaptic markers and show minimal contamination from nuclear (HisH3) or Golgi (GM130) proteins. STX1A and Syb are enriched in SYN fraction but STX1A being an active zone protein is de-enriched from SV fraction (and enriched in SV-S); Syb enriched in SV is used as prey for surface bound capture STX1A. (C) Quantification of SynaptoRed (SR) fluorescence intensity in the SV and SV-S fractions in the supernatant (*input)* and the pelleted material (*pellet*) after centrifugation. SV-S (supernatant during SV-P step ii) shows significantly lower or negligible fluorescence retention than SV (purified vesicle) fraction. Statistical analysis *via* paired t-test. Insets show example tubes indicating pellet visibility post-centrifugation, n = 8, N = 4 from 2 animals. (D) Representative images of SV and SV-S fractions after SynaptoRed dye loading and centrifugation steps. Freshly prepared SV-S samples yield dense, bright pink pellets (*magenta arrows*), which remain visible (but smaller) after two freeze-thaw cycles, confirming loss of vesicle integrity over time.

**Figure 3.**
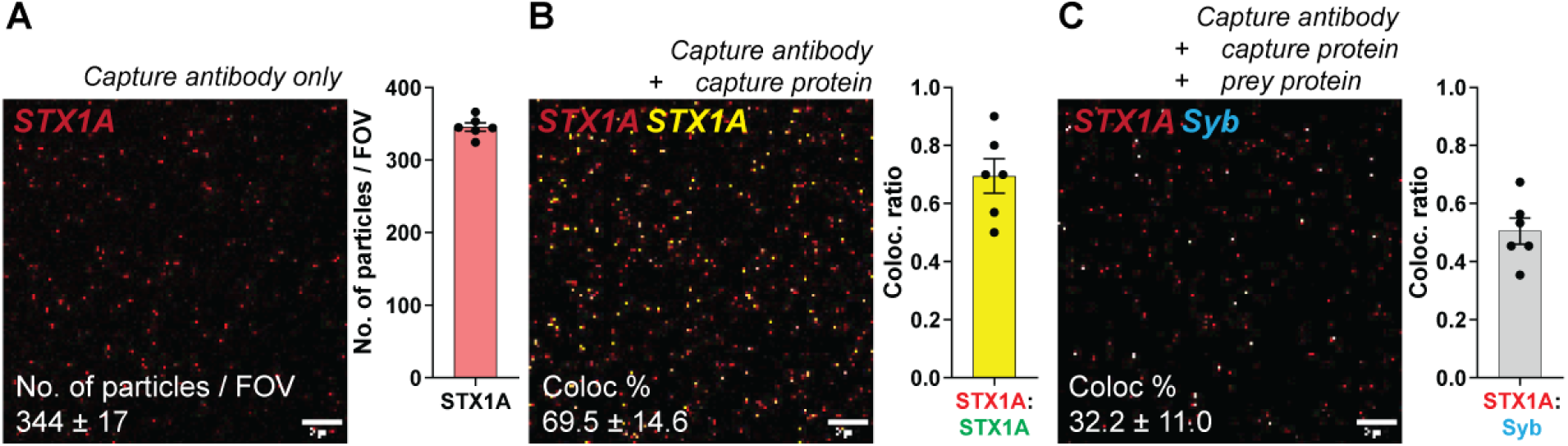
Single-molecule pull-down to probe STX1A-Syb interactions from protein lysates. (A) SIM-Pull assay using capture antibody against syntaxin1 (anti-STX1A) reveals low baseline signal. Representative TIRF image (left) and quantification (right) of functionalized STX1A particles in a FOV. Bar graph shows the number of particles per field (mean ± SEM, 10-12 fields, n = 6, N = 3). Scale bars: 5 µm. (B) Capturing syntaxin-1 proteins from synaptosome homogenate on the STX1A antibody functionalized glass surface. STX1A protein was reprobed using another STX antibody. Dual-color TIRF imaging shows high colocalization (yellow) of red (STX1A capture) and green (STX1A prey) puncta. Quantification of colocalization ratio (mean ± SEM, 10-12 fields, n = 6, N = 3). (C) STX1A-Synaptobrevin (Syb) interaction assay. Capture STX1A protein-antibody complex on functionalized glass surface was incubated with synaptic vesicle lysate (Synaptobrevin as prey). Dual-color TIRF imaging shows colocalization (*white*) between STX1A (*red*) and Syb (*cyan*). Colocalization analysis reveals specific interaction between STX1A and Syb (mean ± SEM, 10-12 fields, n = 6, N = 3), supporting complex formation.

## Preparation of synaptosome homogenate and purified synaptic vesicles (SV)

This protocol primarily describes the isolation of synaptosome-enriched fractions from adult rat forebrains using synaptic protein extraction reagent (Syn-PER™) *via* differential centrifugation. This is a modified protocol optimized in the lab for our experimental use (18–20). The resulting synaptosomes can either be used directly or serve as a source of soluble proteins for synaptosome homogenate (described in step 3) or intact synaptic vesicles (step 4-5) for downstream applications (major step 1 and 2). We recommend preparing the homogenate *via* controlled freeze–thaw cycles and sequential sonication. Also, following mechanical disruption, synaptic vesicles are separated from membrane fragments of the synaptosomes and other debris by ultracentrifugation and purified by a sucrose step gradient [modified from (21,22)]. This protocol yields reproducible, high-quality synaptic proteins and purified intact synaptic vesicles for biochemical and molecular biology assays.

**Timing: [1 day, variable]**

1. Dissect and homogenize rat forebrains
  a. Euthanize an adult rat (8-12 months old) in accordance with institutional animal ethics guidelines.
  b. Dissect the forebrain immediately and place it in ice-cold PBS buffer supplemented with phosphatase and protease inhibitor cocktail. **Critical:** Keep all reagents and tools ice-cold to preserve protein integrity.
  c. Homogenize the forebrain tissue in 2 ml of ice-cold homogenization buffer [0.32 M sucrose, 4 mM HEPES (pH 7.4), supplemented with phosphatase and protease inhibitor cocktail; using a 2-ml glass-Teflon homogenizer. Perform 15–17 gentle strokes.
  d. Rinse the homogenizer with an additional 2 ml of homogenization buffer and combine with the homogenate (total volume: 4 ml). **Note:** In our preparation, we used each hemibrain as technical replicate, making it easier later for balancing the samples during the ultracentrifugation step.
2. Differential centrifugation to enrich synaptosomes (Fig. 2A, *scheme*)
  a. Centrifuge the homogenate at 1,500 × g for 10 min at 4°C to remove cell bodies and debris (pellet P1). **Note:** We recommend setting aside a small volume of each fraction to examine the enrichment of synaptic material *via* ELISA (Fig. 2B) or WB.
  b. Transfer the supernatant (S1) to a clean tube and centrifuge the S1 fraction at 16,100 × g for 20 min at 4°C to pellet synaptoneurosome-enriched fraction (P2). **Critical:** Carefully avoid disturbing the pellet or aspirating the pellet into the S1 fraction.
  c. Resuspend the P2 pellet in 1 ml of homogenization buffer using gentle pipetting.
  d. Prepare a two-step discontinuous sucrose gradient in an ultracentrifuge tube:
    i. Add 4.5 ml of 1.2 M sucrose solution (with 4 mM HEPES supplemented with phosphatase and protease inhibitor cocktail).
    ii. Carefully overlay 4 ml of 0.8 M sucrose solution (with 4 mM HEPES supplemented with phosphatase and protease inhibitor cocktail) on top.
    iii. Layer the resuspended P2 fraction gently on top of the 0.8 M sucrose layer.
    iv. Ultracentrifuge the gradient at 40,000 × g for 70 min at 4°C using a swing-bucket rotor (acceleration and deceleration set to 3). **Critical:** Ensure proper balance of tubes and careful pipetting to maintain gradient integrity.
  e. Using a syringe, carefully collect the synaptosome-enriched fraction at the interface of 0.8 M and 1.2 M sucrose through the tube wall. Volume of collected SYN fraction varies between 0.7 -1 ml, depending on the thickness of the band. Transfer the enriched synaptosomes (SYN fraction) to a clean tube. **Critical:** Avoid collecting material from the top layer to minimize myelin contamination.
  f. We recommend doing a protein estimation so you would have an estimate of protein content in the isolated material.
  g. You can divide the SYN fraction into two parts, aliquot one of each into smaller volumes (50-100 ul) for preparation of homogenates (<u>stopping point</u>). This aids in avoiding repeated freeze-thawing of the material. Additionally, you could use individual fractions to test different lysis buffers depending on the applicability for your experiment.
  h. For the isolation of synaptic vesicles, place the remaining half of each SYN fraction on ice and continue with preparatory step 3. **Critical:** It can be a long day, but we highly recommend isolating synaptic vesicles from the SYN fraction on the same day. Alternatively, it is also possible to store the SYN fraction in Syn-PER buffer (1:1 dilution) at 4°C overnight and proceed with the vesicle isolation the next day (we have not done this).
3. Prepare homogenate from synaptosome-enriched fraction (Fig. 2A, *scheme*)
  a. We prepared homogenate by two methods, i. freeze thaw cycles ii. sonication.
  b. Resuspend the freeze-thawed SYN fraction in 1:2 dilution of ice-cold lysis buffer composition [50 mM Tris-HCl (pH 7.4), 500 mM NaCl, 1% NP-40, 0.5% sodium deoxycholate, 0.1% SDS, 2 mM EDTA, 1 mM DTT, supplemented with phosphatase and protease inhibitor cocktail]. **Note:** Depending on the proteins of your interest and its binding partners, you could additionally use CHAPS (0.5–1%) or digitonin (0.1–0.5%). Furthermore, to break protein–protein complexes more aggressively, you can increase NaCl (up to 500 mM) and/or include 0.5% SDS, as indicated in literature.
  c. Sonicate on ice (3 pulses of 10 seconds each, with 30-second intervals) to rupture the membranes and release proteins. **Critical:** Keep the sample ice-cold throughout; avoid frothing or foam formation.
  d. Centrifuge the sonicated lysate at 21,000 × g for 20-30 min at 4°C to remove large membrane fragments, smaller vesicles and debris.
  e. Carefully collect the supernatant, which contains free floating proteins. Please avoid bubble formation.
  f. Determine protein concentration and prepare single use aliquots. These can be stored at -20°C for a year or more.
4. Isolate synaptic vesicle pellet (SV-P) (Fig. 2A, *scheme*)
  a. Mechanically disrupt isolated synaptosomes using a #23 needle attached to a 1ml syringe by applying 21 strokes. This mixture is further subjected to mild sonication (1-2 pulses of 2 seconds each, with 30-second intervals) on ice. **Critical:** All steps need to be performed on ice or at 4°C. Care should be taken to prevent excessive sonication, as this could disrupt synaptic vesicles.
  b. Centrifuge the sonicated material at 21,000 g for 10 min at 4°C to get rid of larger particles, other vesicular components and membrane debris.
  c. Re-centrifuge the supernatant at 21,000 g for 20 min at 4°C to pellet down synaptic vesicle-pellet (SV-P) fraction, wherein the soluble proteins or dissociated vesicles remain in the supernatant. **Note:** For an initial trial run, you can also use the SV-P to see if you detect some recruitment of the vesicles. It is highly recommended to use the purified SV fraction to obtain optimal results.
5. Purification of synaptic vesicles (SV) (Fig. 2A, *scheme*)
  a. Resuspend the vesicle pellet in 0.5–1 mL of 0.32 M sucrose + 4 mM HEPES, pH 7.4.
  b. Prepare a discontinuous sucrose gradient in ultracentrifuge tubes:
    i. Bottom layer: 1.2 M sucrose (3 mL)
    ii. Middle layer: 0.8 M sucrose (5 mL)
    iii. Top layer: 0.32 M sucrose containing the resuspended pellet (1 mL) **Note:** If you have the smaller ultracentrifuge tubes and the respective swinging rotor, you can use smaller volumes for this step.
  c. Centrifuge the gradient at 100,000 × g for 70 min at 4°C in a swinging-bucket rotor (acceleration and deceleration set to 3).
  d. After centrifugation, collect the synaptic vesicle-enriched band located at the interface between 0.8 M and 1.2 M sucrose using a needle and syringe. The volume of this fraction varies between 0.3 -0.5 ml.
  e. Dilute the recovered vesicle fraction with 3 volumes of Syn-PER buffer to reduce sucrose concentration.
  f. Centrifuge again at 37,000 × g for 45 min at 4°C to pellet vesicles.
  g. Carefully resuspend the final pellet (containing purified synaptic vesicles) in a minimal volume (e.g., 100-200 µL) of storage buffer (Syn-PER buffer with 0.1% DMSO and 0.1% glycerol). **Note:** Synaptic vesicles resolved in storage buffer can be stored at –20°C in small aliquots (about 25 µL) for independent experimental use. You can set aside a small volume for downstream analysis using ELISA (Fig. 2B) or WB assays. **Critical:** Once you thaw an SV aliquot, proceed with labelling and the assay as soon as possible. Once thawed, SV starts to rupture and lose integrity over time, so we prefer using the samples only after one free-thaw cycle (Fig. 2D).

## Key resources table

| REAGENT or RESOURCE | SOURCE | IDENTIFIER |
| --- | --- | --- |
| **Primary antibodies** |  |  |
| Anti-STX1A (rabbit polyclonal) IgG (Clone: 230470F5) | ProteinTech (#83159-6-RR) | RRID: AB_3670852 |
| Anti-RIM1/2 (rat monoclonal) IgG2a (κ light chain) (Clone: SY-53E12) | Synaptic Systems (#140217) | RRID:AB_2924949 |
| RIM1 antibody PDZ domain (rabbit polyclonal) | Synaptic Systems (#140003) | RRID:AB_887774 |
| Anti-Syb1 (Guinea pig polyclonal) | Synaptic systems (#104004) | RRID:AB_2619755 |
| Anti-VAMP-1/2/3 (Guinea pig polyclonal) | Synaptic systems (#104104) | RRID:AB_2619756 |
| Anti-Rab 3A (rabbit polyclonal) (clone: K-15) | Santacruz Biotechnology (#sc308) |  |
| Anti-STX1A/1B (mouse monoclonal) (clone: 2D3B5) | ProteinTech (#66437-1-Ig) | RRID:AB_2881807 |
| Anti-RIM1/2 (Guinea pig polyclonal) | Synaptic Systems (#140005) | RRID:AB_2661872 |
| Anti-SNAP25 antibody (Guinea pig polyclonal) | Synaptic Systems (#111004) | RRID:AB_2192329 |
| **Secondary fluorescent antibodies** |  |  |
| Goat anti-rabbit IgG A488 | Abcam (#ab150077) |  |
| Goat anti-rabbit IgG (H+L) Highly Cross-Adsorbed Secondary Antibody, Alexa Fluor™ 568 | Invitrogen (#A11036) |  |
| Goat anti-rabbit IgG (H+L) Highly Cross-Adsorbed Secondary Antibody, Alexa Fluor™ 647 | Invitrogen (#A21245) |  |
| Goat anti-mouse IgG (H+L) Highly Cross-Adsorbed Secondary Antibody, Alexa Fluor™ 647 | Invitrogen (#A21246) |  |
| Goat anti-mouse IgG A488 | Abcam (#ab150113) |  |
| Goat anti-mouse IgG A647 | Abcam (ab150115) |  |
| Goat anti-mouse IgG 405 | Sigma Aldrick (#50185-1ML-F) |  |
| Goat anti-guinea Pig IgG (H+L) Highly Cross-Adsorbed Secondary Antibody, Alexa Fluor™ 647 | Invitrogen (#A21450) |  |
| Goat anti-guinea Pig IgG A488 | Abcam (#ab150185) |  |
| Goat anti-guinea Pig IgG A647 | Abcam (ab150187) |  |
| Goat anti-Rat IgG (H+L) Cross-Adsorbed Secondary Antibody, Alexa Fluor™ 647 | ThermoFischer Scientific (#A-21247) |  |
| Donkey anti-Rat IgG (H+L) Highly Cross-Adsorbed Secondary Antibody, Alexa Fluor™ 488 | Invitrogen (#A21208) |  |
| Streptavidin, Alexa Fluor™ 488 conjugate | Biolegend (#405235) |  |
| Streptavidin, Alexa Fluor™ 647 conjugate | Biolegend (#405237) |  |
| **Secondary Li-COR antibodies** |  |  |
| IRDye 680 RD Goat anti Mouse | LI-COR Biosciences (#926-32210) |  |
| IRDye 800CW Goat anti Mouse | LI-COR Biosciences (#926-68070) |  |
| IRDye 680 RD Goat anti Rabbit | LI-COR Biosciences (#926-68071) |  |
| IRDye 800CW Goat anti Rabbit | LI-COR Biosciences (#926-32211) |  |
| **Secondary HRP-conjugated antibodies** |  |  |
| Goat anti-Mouse IgG (H+L) Secondary Antibody, HRP | ThermoFischer Scientific (#32430) |  |
| Goat anti-Rabbit IgG (H+L) Secondary Antibody, HRP | ThermoFischer Scientific (#31460) |  |
| Goat anti-Guinea Pig IgG (H+L) Secondary Antibody, HRP | ThermoFischer Scientific (#A18769) |  |
| Goat anti-Rat IgG (H+L) Secondary Antibody, HRP | ThermoFischer Scientific (#31470) |  |
| **Chemicals** |  |  |
| Buffer, PBS (10X, Dulbecco's) powder pack | VWR (#A0965.9010) |  |
| EDTA | CarlRoth (#1E23.1) |  |
| Proteinase K, recombinant, PCR grade | ThermoFischer Scientific (#EO0492) |  |
| (3-Aminopropyl)triethoxysilane APTES, 98% | ThermoFischer Scientific (#A10668.22) |  |
| Acetone ≥99% | VWR-Avantor (#VWRC20063.365) |  |
| Pierce™ Hydroxylamine-HCl | ThermoFischer Scientific (#26103) |  |
| Pierce™ SATA (N-succinimidyl S-acetylthioacetate) | ThermoFischer Scientific (#26102) |  |
| Sulfo-SMCC sodium | BioConnect (#HY-D0975) |  |
| Pierce™ Dextran Desalting Columns (5K MWCO) | ThermoFischer Scientific (#43233) |  |
| Dispensing tips | VWR (#613-2062) |  |
| DTT (dithiothreitol) | ThermoFischer scientific (#R0861) |  |
| β-ME (β-mercaptoethanol) | Gibco (#21985023) |  |
| Synaptic Protein Extraction Reagent (Syn-PER™) | ThermoFischer Scientific (#87793) |  |
| Protease Inhibitor Cocktail Set III, EDTA-Free - Calbiochem | Sigma Aldrich (#539134-1ML) |  |
| PhosSTOP™ | Sigma-Roche (#4906845001) |  |
| HEPES | Sigma Aldrich (#H4034) |  |
| Sucrose | Sigma Aldrich (#S0389) |  |
| DMSO | Sigma Aldrich (#20-139) |  |
| Glycerol | VWR-Avantor (#7044.1000) |  |
| Tris-HCl | ThermoFischer Scientific (#228030010) |  |
| NaCl | Carl Roth (#3957.2) |  |
| NP-40 | bioPLUS (#22040045) |  |
| Sodium deoxycholate | Sigma Aldrich (#D5670) |  |
| SDS (pellet form) | Carl Roth (#CN30.3) |  |
| SynaptoRed C2 (lipophilic dye, equivalent to FM®4-64) | Biotium (##70027) |  |
| Rhodamine B-DHPE | Thermo Fisher Scientific (#R7481) |  |
| Bodipy FL C5-HPC | Invitrogen (#D3922) |  |
| Trypsin (cell culture grade) | Roche (#03358658103) and Gibco (#15400-054) |  |
| PMSF (Phenylmethylsulfonyl fluoride) | Sigma-Aldrich (#P7626) |  |
| Bovine Serum Albumin (BSA) Fraction V, NZ-Origin | Carlroth (#8076,4) |  |
| Gibco™ Horse Serum, New Zealand origin | ThermoFischer Scientific (#16050130) |  |
| Normal Goat Serum | Abcam (#ab7481) |  |
| 1-Step™ TMB ELISA Substrate Solutions | ThermoFischer Scientific (#34028) |  |
| Sulphuric acid (H<sub>2</sub>SO<sub>4</sub>) | Avantor (#7664-93-9) |  |
| **Commercial Assays** |  |  |
| BCA protein assay kit | ThermoFisher Scientific (#23227) |  |
| ELAST ELISA Amplification System | Revvity (#NEP116E001EA) |  |

| **Softwares** |  |  |
| --- | --- | --- |
| Image Studio/ Image Studio Lite | LiCOR Bio |  |
| FIJI (ImageJ) | NIH | https://fiji.sc,<br>RRID:<br>SCR_002285 |
| GraphPad Prism 10 | PRISM Academy | https://www.graphpad.com,<br>RRID:<br>SCR_002798 |
| **Equipment and instruments** |  |  |
| Ultracentrifuge (for synaptosome isolation and SV pelleting) | Beckman Coulter Model Optima-90 (SN CXE14C01) Rotor SW41Ti and SW45Ti |  |
| Benchtop Ultracentrifuge (SV purification and labelling) | ThermoScientific X4R Pro |  |
| Shaker | Linear: Laboshaker;<br>Orbital: Rotamax |  |
| Centrifuge | Eppendorf 5804R,<br>Heraeus-Fresco21 |  |
| High-precision glass coverslips (No. 1.5H, 25 mm) | VWR (#631-1584) |  |
| Thermomixer | Eppendorf (#5382000015) |  |
| Plate reader | Tecan Model Spark M10 |  |
| Plasma cleaner (optional) | Diener Femto |  |
| Quick release magnetic and non-magnetic imaging chambers | Warner Instruments (#QR-40LP) |  |
| Culture dish microincubator | Warner instruments (#DH-40iL) |  |
| Fusion 100-X Touch infusion syringe pump | KR Analytics |  |
| BD Plastipak Syringes Luer Lock | BD Syringes (#309630 and #309628) |  |
| Fittings and tubing | IDEX/Darvin Microfluidics (#CIL-P-642, #CIL-P-857, #CIL-P-678, #BL-PTFE-1608-20, #CIL-XP-245X, #CIL-P-632, #CIL-P-634, #CIL-P-603, #CIL-P-702, #CIL-P-250X, #CIL-P-259X) |  |
| TIRF microscope | Nikon live-cell TIRF and epifluorescence microscope |  |

## Materials and equipment setup

- All surface functionalization steps should be performed in a clean, dust-free environment using powder-free gloves. Optionally you could use acid-washed or plasma-cleaned coverslips.
- Use of a humidified chamber during incubation steps to prevent drying of reagents on coverslips, when using low volumes for antibody, protein incubation steps.
- Thermomixer setup for vesicle labelling (temperature set to 25°C and shaking speed to 300–400 rpm)
- It is recommended to use low-retention Eppendorf tubes to minimize protein/vesicle/dye loss.
- TIRF microscope equipped with 100x oil objective (NA 1.49 or higher) for optimal resolution of single vesicle events.
- Laser power and exposure should be adjusted to minimize photobleaching while ensuring adequate signal-to-noise ratio (SNR).
- FIJI (ImageJ) macros may be customized for automated vesicle detection and background subtraction.

### Buffer/solution Recipes

**Note:** We recommend filtering the buffers using a 0.22 µm filter (indicated in the respective recipes). This would aid in getting rid of any small dust/particles etc. which could result in unwanted artefacts during imaging. Autoclaving water before use is not necessary.

- Silanization buffer [2% (v/v) 3-Aminopropyltriethoxysilane (APTES) in dry acetone]
  - It is recommended to use drying beads to get rid of moisture from acetone, optionally purging with inert nitrogen-N_2_ gas would also do the trick.
  - Buffer should be prepared fresh and used within 30 min max. ***Caution!*** *APTES* is corrosive and may cause severe skin, eye, and respiratory irritation. Handle inside a certified chemical fume hood with a lab coat, nitrile gloves, and protective eyewear. Avoid exposure to air/moisture prior to use to prevent hydrolysis and unwanted polymerization. Store tightly sealed under inert gas or in a desiccator. Dispose of spills and unused material as hazardous organic waste as per institute regulations. ***Caution!*** *Acetone* is highly flammable and volatile; please keep away from ignition sources. Use only in a fume hood. Avoid prolonged skin contact to prevent defatting and irritation. Collect all acetone waste in designated flammable liquid waste containers as per institute regulations.
- Phosphate buffer [1x PBS (137 mM NaCl, 2.7 mM KCl, 10 mM Na₂HPO₄, 1.8 mM KH₂PO₄) and 10 mM EDTA, adjusted to pH 7.4]
  - Prepare with ultrapure water and adjust pH to 7.4 before adding EDTA. EDTA should be added after pH adjustment to prevent chelation interference during pH adjustment.
  - Filter-sterilize using a 0.22 µm filter and store at 4°C for up to 6 months.
  - We recommend preparing a 5x solution and aliquoting into ready-to-use parts for experiments. Aliquots can be stored at 4°C for up to 6 months.
- Homogenization buffer [0.32 M sucrose, 4 mM HEPES (pH 7.4), phosphatase inhibitor (1:10000), EDTA-free protease inhibitor cocktail (1:1000)].

- Prepare a stock solution of 2 M HEPES in ultrapure water and adjust pH 7.4. The solution is stable at RT for 6 months to 1 year. If the solutions appear cloudy or slightly translucent, please discard immediately. You can also use this solution to prepare solutions for the sucrose gradient, which could also be prepared in a big batch and stored indefinitely as a single time ready to use aliquots.
- Weigh and add the required amount of sucrose to diluted 4 mM HEPES buffer. You can prepare a larger batch and store indefinitely as a single-time ready-to-use aliquots at - 20°C.
- Phosphatase and protease inhibitor cocktail should be added immediately before use.
- Lysis buffer [50 mM Tris-HCl (pH 7.4), 500 mM NaCl, 0.5-1% NP-40, 0.5% sodium deoxycholate, 0.1% SDS, 2 mM EDTA, 1 mM DTT, supplemented with phosphatase and protease inhibitor cocktails]
  - Prepare stock solutions (i) 1 M Tris-HCl in ultrapure water and adjust pH 7.4; (ii) 10 % SDS (label the bottle/tube with warning signs) and (iii) 5% sodium deoxycholate solution in ultrapure water. All stock solutions are stable at RT for 6 months to 1 year. If the solutions appear cloudy or slightly translucent, please discard immediately.
  - You can also prepare a 10 mM DTT stock solution and store ready to use aliquots at - 20°C. These aliquots could be stored indefinitely.
  - Weigh NaCl and EDTA and dissolve in 50 mM Tris buffer prepared from the stock solution.
  - Add respective volumes from SDS (1:100), sodium deoxycholate (1:10), DTT (1:10) stock solutions as well as NP-40 (0.5-1%). This solution is stable at RT for a few months. You could also store it at 4°C to increase its shelf life but note that sometimes you can see a bit of turbidity due to SDS precipitation which should clear up as SDS dissolves when vortexed.
  - Finally add phosphatase and protease inhibitors prior to use. Buffer supplemented with inhibitors can be stored at -20°C in single time use aliquots. We do not recommend using repeated freeze-thawed buffer aliquots.
- Blocking buffer [2.5% (w/v) BSA and 2.5% normal horse serum in phosphate buffer]
  - Weigh the required amount of BSA and dissolve in phosphate buffer. Add the required volume of horse serum to make up to 4% of the solution.
  - Filter the solution using a 0.22 µm filter and store at 4°C. Use within a week.
  - You can also prepare a big batch and freeze ready to use pre-filtered aliquots at -20°C for a year.
  - We recommend using the same batch of buffers for imaging all the conditions of a particular set of experiments.
- Syn-PER buffer [Syn-PER reagent supplemented with 1x phosphatase and 1x protease inhibitor cocktail]
  - Add inhibitors to the reagent and filter the solution using a 0.22 µm filter and store at 4°C. Use on the same day. **Note:** If preparing larger volumes, we recommend freezing ready to use aliquots. Do not use solutions after 2 freeze thaw cycles.
  - To prepare Syn-PER storage solution (add 0.1% DMSO and 0.1% glycerol) to store intact synaptic vesicles at -20°C.
- Imaging solution [1:1 mixture of blocking buffer and Syn-PER solution]

- Prepare and use as required. Discard excess solution at the end of the day.

## Pull-down of proteins from synaptosome homogenate

This major step enables the selective immobilization and detection of capture/target proteins from a synaptosome homogenate onto a functionalized glass surface. First, non-specific binding sites are blocked and incubated with the protein sample; secondly by using specific detection antibodies, this protocol helps isolate and visualize proteins directly from native cellular material. The use of fluorescence-based detection allows for quantitative assessment of protein binding efficiency and comparison across experimental conditions, validating the efficacy of the surface functionalization and pull-down strategy.

This step could be further explored to examine conformations, aggregation and the specific interaction/pull down with prey proteins specific to the immobilized capture protein(6,7).

**Timing: [5-6 h]**

1. Blocking of reactive sites
  a. Wash the functionalized coverslips (in a 6-well plate) with 1.5 ml of phosphate buffer twice (quick wash). Remove the solution completely.
  b. Incubate with 1.5 or 2 ml of blocking solution (2.5% BSA and 2.5% NHS in phosphate buffer), mild agitation for 1 h at RT. **Note:** Blocking is essential, the time of incubation could be reduced to 30 min. Agitation is not important. **Critical:** Store one coverslip after this step to measure background fluorescence (This will be your antigen-delete and secondary antibody-delete control).
2. Loading your capture protein
  a. Thaw the vial containing synaptosome homogenate on ice and dilute it in blocking buffer. Prepare an appropriate dilution based on the protein concentration determined by LOD assays.
  b. The volume of liquid should be sufficient to cover the entire surface of the coverslip, incubate for 1-2 h at RT. Please do not forget to prepare coverslips without the antigen solution as your antigen-delete control, you can incubate these with just the blocking buffer. **Note:** In case of lower protein concentration of your prepared material or to increase the efficiency of pull-down, you can place a 150 μl drop of protein sample on a parafilm strip and invert the coverslip (functionalized surface facing the liquid downward). Additionally, in case of sensitive proteins, you can also choose to incubate the protein homogenate overnight at 4°C.
  c. Remove the unbound material and wash the coverslips with phosphate buffer (2 x quick washes and 2 x washes with 5 min incubation, with agitation).
  d. Perform a second blocking step for at least 30 min, followed by washing steps as described in step 2c.
  e. Coverslips containing the capture protein-antibody complexes can be stored in phosphate buffer at 4°C in dark (supplemented with sodium azide for longer durations). We always proceeded with the next major step within a week.
3. Detecting loading efficiency
  a. Incubate the coverslip with a detection antibody (respective dilution as determined) in blocking buffer for 1 h. Repeat washing steps (described in 2c.). **Note:** Dilutions for primary and secondary antibodies for ICC steps are always in the range of 1:500 to 1:1000 (depending on the specificity and sensitivity of your antibodies in use.
  b. Incubate the coverslip with secondary antibodies of your choice (different colors for each, capture and detection, respectively), for 1 h at RT. **Note:** If you are using a fluorophore-conjugated detection antibody in step 3a., then only add the secondary antibody for your capture antibody. You can also merge the two steps together. **Critical:** Use one set of coverslips to examine the efficiency of binding of your capture protein to the capture antibody. A colocalization analysis of the two fluorophores will yield the exact ratio of bound versus unbound sites of the capture antibody/protein complex. These values are important for determining the functionality of your antibody post modification step.
  c. You can also just add the secondary antibody to a functionalized coverslip without the antigen to know the total number of functional antibodies on your surface. **Critical:** The surface density of about 300-400 antibody molecules per field of view-FOV (please see Fig, S3). For a good spatial separation and resolution these particles should cover not more than 20 % of the FOV. **Note:** If the values from this experiment and step 2 differ quite significantly, you would need to retrace back the steps and use a different antibody either capture or detection due to difference in reactivity of epitope bindings, reducing the effective detection. Also, we have observed that the pull down of capture protein *via* the functionalized antibody is in the range of 60-80 % when detected with a second antibody specific to the capture protein. It is important to note here that the orientation of the antibody during functionalization is not controlled and thus the ease of accessibility to the functional epitope alters this ratio. Similarly, when probing the prey protein post capture-prey interaction, this ratio drops below 50 %.

## Recruitment of synaptic vesicles assay

This step enables the real-time visualization of tethered synaptic vesicle (SV) recruitment to the surface *via* the immobilized capture protein-antibody complexes. By fluorescently labelling isolated SVs and incubating them with pre-functionalized coverslips, this assay helps determine the binding affinity, specificity, and dynamics of vesicle-protein interactions under near-physiological conditions. Careful timing, imaging calibration, and buffer control are essential to maintain vesicle integrity and avoid nonspecific background fluorescence. This protocol facilitates downstream analyses of vesicle recruitment efficiency based on protein-protein binding kinetics.

**Timing: [4-6 h, variable]**

1. Labelling synaptic vesicles with fluorescent dyes
  a. Thaw an aliquot of SV on ice (usually 10-15 min), and add 9 parts of ice-cold imaging buffer [1:1 of Syn-PER and blocking buffer]
  b. To this mix, add the SynaptoRed dye (1:10000 final dilution). Incubate for 30 min at RT using mild agitation (Thermomixer -200-300 rpm).
  c. Make up the volume to 0.8-1 ml using ice-cold imaging buffer and centrifuge the SV at 37,000 g for 30 min at 4°C. **Note:** You will be able to see a clear pink pellet at the bottom of the tube (Fig. 2C-D).
  d. Discard the supernatant containing the excess dye. Without disturbing the pellet, add 300 μL of ice-cold imaging solution and discard the supernatant.
  e. Resuspend the pellet carefully in 200 μl of ice-cold imaging solution and store the tube on ice. **Critical:** It is essential that this step be performed immediately before the imaging experiments (described in the next steps). Any residual SR loaded-SV material should be discarded at the end of the day. **Note:** In this protocol, we have successfully used SynaptoRed (bright fluorescence and easy to work with). Other lipid dyes like rhodamine, Bodipy, have also worked comparatively well.
2. Tethering of synaptic vesicle *via* capture-prey protein binding
  a. Assemble the coverslips containing the capture protein-antibody complex loaded on the imaging chamber and apply 400-500 μl of imaging solution to the coverslips. **Note:** Before incubating the SR-SV material, it is recommended to acquire 10-15 images (coverslip-specific background correction in later steps). Also, for all imaging conditions optimize the volume of buffer that gives you the best resolution and keep the volume constant for the remainder of the experimental conditions.
  b. Dilute the SV-SR material in imaging buffer, we used a 1:200/300 dilution, when starting with a 20-25 μg protein concentration in an aliquot.
  c. Remove the imaging buffer from the mounted coverslip and load the SR-SV mixture, incubate for 30-45 min at RT. **Critical:** Excess SV concentration and longer incubation times can lead to bursting of vesicles and ultimately result in higher background fluorescence (Fig. S5 and S6A). **Note:** Since the complete set-up is on the objective, you can image every 10-15 min in TIRF mode to check for the SR fluorescence (bound to the surface). Post-incubation, flow through imaging buffer for about 1 min (flow rate 2 ml/min, 4-5 changes of complete solution within imaging chamber). Let it stand for 2 min, repeat the flow again for 30 sec. This step ensures the riddance of all sterically bound SVs (Fig. S5).
3. Imaging vesicle recruitment events
  a. Start TIRF imaging and acquire 10–15 fields of view (FOVs) per coverslip. Each FOV should span at least 50–100 μm² or include about 100 particles. Our imaging parameters were set to detect 300+ antibody particles/FOV. This was considered optimal as we only had ∼ 70% colocalization with capture-capture antibody and ∼50% capture-prey antibody complex (Fig. 3)
  b. Image acquisition should be performed under minimal laser power exposure to prevent dye bleaching. **Note:** Due to time constraints on imaging as many parameters as possible on the same day, it is optional to save the images in processed formats (e.g., background-subtracted, contrast-adjusted). This can also be done later using an image processing software (e.g. Fiji ImageJ).
  c. Ideal images should look as described in (Fig. 4A and Fig. S5). Do not forget to save the images after acquisition. Also, saving the raw files which contain the meta data of your acquired images, might be beneficial for setting your analysis parameters later. Imaging parameters for the images depicted in the manuscript 66.56 x 66.56 microns, 1952 x 1952 pixels, 29.32 pixels/micron; 16-bit images.
4. Control experiment: Antibody blocking of prey proteins on SVs This protocol allows you to determine the extent of vesicle recruitment inhibition when prey proteins are masked. A decrease in SV binding in the antibody-treated groups (relative to untreated) confirms the specific involvement of prey proteins (Syb/SNAREs and/or Rab3a) in recruitment *via* capture proteins (STX1A and/or RIMs, respectively). The dilution series helps assess the saturation threshold of antibody blocking, and partially blocked SVs may reveal cooperative or multivalent interactions between the vesicle and capture surface.

a. Add antibodies against prey proteins at 1:1, 1:2, and 1:3 dilutions (relative to standard working concentration and determined LOD values, Fig. 1 and S1) and incubate for 15-30 min at RT. **Note:** Prepare one reaction per antibody dilution.
b. Repeat steps described earlier 2a-e. Blocking of interacting epitopes would result in lower tethering of synaptic vesicles, please see (Fig. 4B). **Note:** Excess antibodies in this step would then be significantly diluted and washed off in the next steps, thus there is no need to specifically pellet down the synaptic vesicles.
5. Control experiment: Digestion of SV surface proteins If recruitment is due to specific protein–protein interactions (STX1A-SNAREs and RIM1-Rab3a, respectively), then disrupting SV surface proteins should reduce or abolish binding, while non-specific interactions (e.g., lipid-lipid or charge-based) would be less affected.
  a. Thawed SV aliquots should be diluted in Syn-PER buffer (Reaction needs to be carried out in the absence of serum/BSA, as proteins would inhibit trypsin).
  b. Add trypsin (e.g., 5 µg/ml) to the SV preparation.
  c. Incubate at 37°C for 10-15 min, with gentle agitation (Thermomixer, 37°C, 200-300 rpm) **Critical:** Shorter incubations preserve vesicle integrity; adjust based on pilot titration. Longer incubation times (>30 min) significant loss of synaptic vesicle integrity was observed, as determined by the decrease in pellet size).
  d. Immediately inhibit trypsin by adding PMSF (1 mM) and 2% NHS in the imaging buffer. **Note:** It is optional to use a centrifuge (as described in 1c., discard supernatant and wash the pellet once with 300 µl of ice-cold imaging solution.) You can also directly proceed with loading the vesicles with SR dye (in blocking buffer), excess of the reagents would be washed away subsequently.
  e. Load these synaptic vesicles with SR dye by following steps 1 and 2 as described earlier. Proteolytic cleavage of interacting epitopes would result in lower tethering of synaptic vesicles, please see (Fig. 4B). **Note:** You can also use other molecular cutters specific to your protein of interest, please optimize the incubation time accordingly.

**Figure 4.**
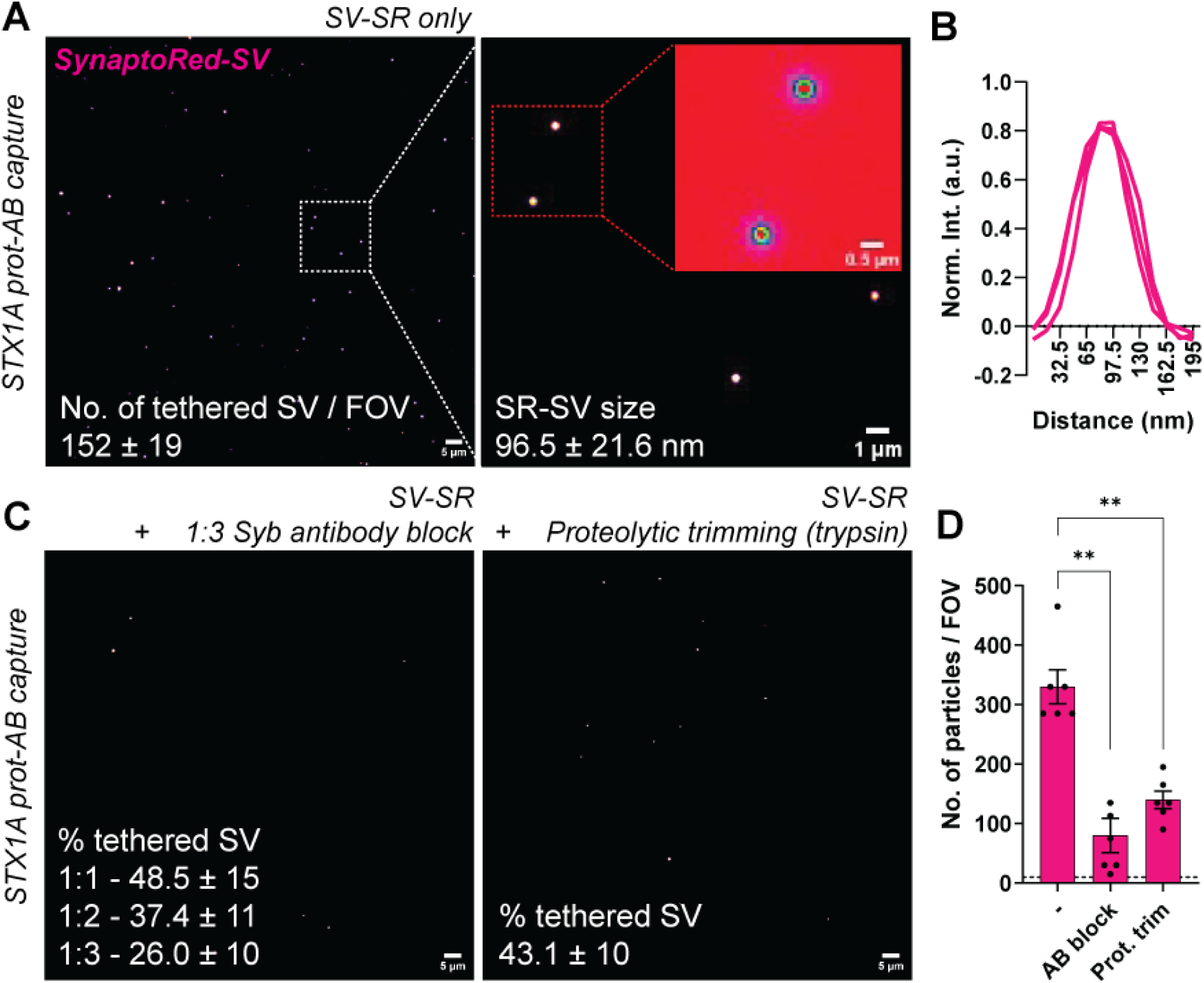
Synaptic vesicle recruitment *via* capture-prey (STX1A-Syb) system. (A) Representative TIRF images depicting recruitment of synaptic vesicles loaded with SynaptoRed (SR-SV) using extended SIM-Pull assay protocol *via* the STX1A-Syb capture-prey system. Scale bars: 5 µm, 1 µm and 0.5 µm, respectively. (B) Line plots depicting the average spot profile or signal intensities of tethered SR-SV to the surface. (n = 4). Quantification is representative of 2 independent experiments. (C) Representative images show that blocking prey protein (Syb) *via* antibody blocking, or proteolytic degradation of the binding epitope leads to decrease in recruitment of synaptic vesicles. (D) Quantification of number of particles per FOV (mean ± SEM, 10-12 fields, n = 6, N = 3). One-way ANOVA, * *p* < 0.05, ** *p* < 0.01.

## Live imaging of protein–protein conjugation on recruited synaptic vesicles

This step enables the visualization of surface-accessible or interacting proteins on intact, recruited synaptic vesicles under near-physiological conditions using live immunocytochemistry with fluorophore-conjugated secondary antibodies. The procedure avoids fixation to preserve integrity of the dye incorporated vesicles.

**Note:** This step is completely optional. We only performed this step once we had completely optimized the protocol for synaptic vesicle recruitment. We recommend this step to indeed show that the combination of prey and capture proteins are involved in the tethering the SV to the surface.

**Timing: [1-2 h]**

1. Labelling capture and prey protein complex *via* ICC
  a. Dilute primary antibodies against the prey protein to be detected in imaging buffer (dilution/concentration specific to your experiment) and incubate coverslips for 30 min - 1 h at RT in dark, protected from light. **Critical:** It is not recommended to do this step during the incubation of SVs, since this would either block the epitopes of the capture or prey proteins and significantly affect the tethering of synaptic vesicles, resulting in lower readout/false negatives. **Note:** We recommend acquiring 10-15 images before starting this protocol, as frequent incubation and washing steps result in a loss of tethered SV (Fig. 5C).
  b. Gently wash the coverslips with imaging buffer to remove unbound antibodies (2 x quick washes and 2 x with 5 min incubation. Ensure efficient washing to get rid of excess antibodies.
  c. Dilute secondary antibodies against the primary antibody species (both prey and capture) in imaging buffer (again, dilution/concentration specific to your experiment and choice of antibodies) and incubate coverslips for 30 min - 1 h at RT in dark, protected from light. **Note:** Ensure minimal disturbance during incubation to preserve vesicle integrity and we recommend doing this in the imaging chamber itself. We suggest having multiple imaging chambers, so it’s easier to utilize the time of incubation to image other coverslips/conditions.
  d. Gently wash the coverslips with imaging buffer to remove unbound antibodies (2 x quick washes and 2 x with 5 min incubation.
  e. Immediately proceed to imaging all the three channels. Ideal representative images have been depicted in Fig. 5B and S4. **Note:** SynaptoRed fluorescence decreases over time due to multiple washing steps (dilution of the dye and/or vesicle rupture), You could also lower the volume of solution as much as possible. You could additionally optimize the use of alternative antibodies to tag/identify different proteins or the type of vesicles being recruited *via* this protein-protein interaction.

**Figure 5.**
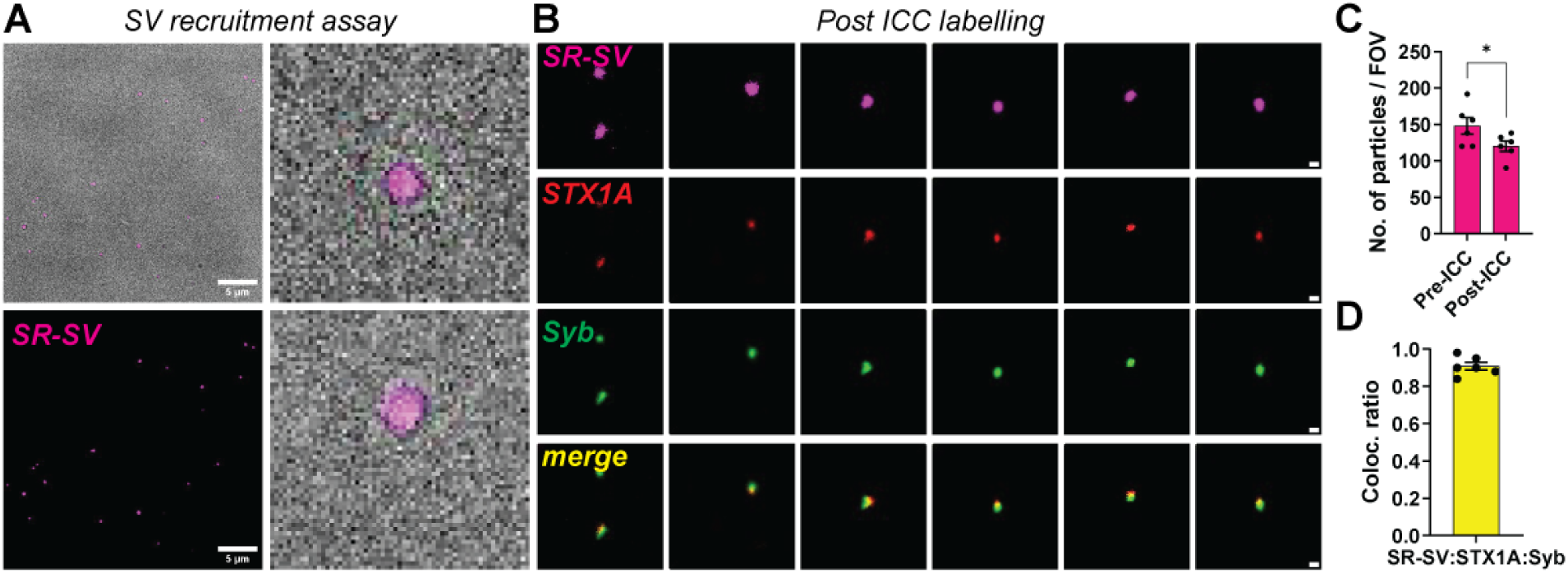
Immunocytochemistry to probe capture-prey (STX1A-Syb) protein complexes tethering SV recruitment. (A) Representative images showing brightfield and SV channels (TIRF mode), respectively; before performing ICC. (B) Zoomed in panels depicting tethering of SV is indeed *via* STX1A and Syb interaction. Zoomed out FOV is depicted in SI, fig. S4A. (C) Bar plots depict quantified number of particles per FOV, wherein we see decrease in tethered SV particles post ICC (mean ± SEM, 10-12 fields, n = 6, N = 3). Students-*t* test, * *p* < 0.05. (D) Colocalization analysis reveals specific interaction between STX1A and Syb supporting complex formation in tethering SV to the surface (mean ± SEM, 10-12 fields, n = 6, N = 3).

## Image analyses and quantification

This step quantifies synaptic vesicle recruitment events to assess capture efficiency and the specificity of protein-protein interactions..

**Timing: [variable]**

1. Image Analysis
  a. Analyze acquired images using ImageJ/Fiji or custom scripts (e.g., MATLAB, Python), depending on your usage/feasibility. We worked with Fiji ImageJ plugins and described the workflow in the SI.
  b. Define a threshold for vesicle detection based on fluorescence intensity and spot size (pixel diameter).
  c. Use particle counting or peak detection plugins to identify individual SVs. A short description is summarized in the SI, section 1B.
  d. Calculate:
    i. Vesicle density (no. of vesicles per FOV or μm², mean ± SD across FOVs), please see Fig. 3A, 4B, 5C and S3.
    ii. Spot intensity profiles (for vesicle size or dye incorporation variability), please see Fig. 4A.
    iii. Coefficient of variation across FOVs and experimental conditions, please see Fig. S3.
  e. Normalize values against blank coverslip or blocking control (surface functionalized (i) without secondary AB or (ii) primary delete with secondary antibody control).
2. Data Interpretation and Statistical Testing
  a. Compare SV binding levels between:
    i. Untreated vs antibody-blocked SVs, please see Fig. 4B.
    ii. Untreated vs trypsin-digested SVs, please see Fig. 4B. **Note**: A >70% reduction in SV density upon antibody blocking or trypsin digestion confirms **specific, protein-mediated vesicle capture**. Residual background (<10%) may arise from charge-based or hydrophobic interactions.
    iii. Additionally, you can customize your assay - wild-type *vs* mutant capture/prey proteins.
  b. Perform statistical analysis:
    i. Use unpaired *t*-test or one-way ANOVA (for ≥3 conditions).
    ii. Represent data in box plots, bar graphs, or scatter plots with appropriate error bars.
    iii. Calculate fold change or percentage inhibition relative to controls.

## Expected outcomes

- Using this protocol, one can expect robust, quantifiable, and reproducible visualization of synaptic vesicle (SV) recruitment to surface-immobilized protein-antibody complexes.
- The methodology described for surface functionalization and antibody coupling yields high-quality coverslips with minimal background fluorescence (<5-10 molecules/FOV).
- Upon incubation with fluorescently labeled SVs, users can expect to observe discrete fluorescent puncta, typically 100-200 vesicles per FOV, corresponding to specific vesicle-tethering events *via* protein-protein (capture-prey) interactions.
- Representative images should display punctate fluorescence localized to the capture surface, with minimal diffuse background signal, please see Fig.4A, 5A and S4.
- The signal intensity and distribution will vary depending on the abundance and affinity of the capture protein, prey protein accessibility, antibody sensitivity and blocking or digestion treatments. Specific recruitment will significantly decrease upon masking of prey proteins *via* antibody blocking or proteolytic digestion, validating the molecular specificity of the interaction, computed as quantitative outcomes (vesicle density per μm² or FOV).
- Importantly, this platform can be extrapolated to detect changes in vesicle recruitment in response to mutations, competitor proteins, or chemical modulators, and can be extended to other vesicle systems such as exosomes, secretory vesicles, or endosomal populations.
- Additionally, dynamics of vesicle association and potential lateral mobility of tagged vesicles could be monitored using time-lapse imaging.

## Background correction, quantification and statistical analysis

- Images acquired *via* TIRF microscopy can be processed using Fiji-ImageJ or equivalent image analysis software. Raw images are first corrected for background fluorescence by subtracting average signal from blank coverslips imaged under identical conditions.
  - You would need a blank imaging buffer channel and a functionalized coverslip without SR-SV to determine:
    - Camera offset (dark image)
    - Coverslip autofluorescence (without any antibody, primary antibody only, secondary antibody only)
    - Non-specific fluorescence in imaging buffer
- Subtract the average background (from 6–10 images) for each coverslip batch. **Critical:** Ensure background levels are <0.02 molecules/μm² (<5-10/FOV) to validate assay sensitivity.
- Particle detection is performed using threshold-based analysis or spot detection plugins (e.g., TrackMate), with vesicle-sized puncta identified based on defined criteria for fluorescence intensity and area (typically 2–5 pixels in diameter, above local background threshold).
- Only fields of view (FOVs) with uniform illumination and no physical artefacts are included in the analysis, please see SI, Fig. S2B-D.
- Vesicle density is calculated as the number of fluorescent puncta per FOV or μm² (please see Fig. 3A, 4B, 5C and S3.) across at least 10–15 FOVs per condition, per experimental replicate. We also recommend doing the antibody modification and surface functionalization at least three times and considering this as an experimental replicate. Please make sure to prepare enough coverslips for each condition to ensure you have enough for all testing conditions + an additional 2.
- Please stop acquiring data if background fluorescence varies more than 10% between the same coverslips of the same preparation, or if vesicle aggregation or lysis is observed, please see Fig.S3-S6.
- Statistical comparisons between conditions (e.g., untreated vs. antibody-blocked, or trypsinised, or between native and mutant variants) are conducted using appropriate statistical analyses. Data should be reported as average of experimental replicates ± standard deviation or standard error of mean, with a significantly assigned statistical test.

## Limitations

While this protocol enables sensitive and specific visualization of synaptic vesicle (SV) recruitment *via* protein–protein interactions, several limitations should be carefully considered.

- The functionalization of glass coverslips depends on consistent and uniform thiolation and cross-linking efficiency; minor deviations in pH, incubation time, or reagent freshness can impact surface reactivity and lead to uneven antibody coverage or increased background fluorescence.
- Also, the orientation of antibody functionalization across the glass coverslip cannot be controlled
- The use of isolated native SVs, although advantageous for preserving physiological topology and composition, introduces variability across preparations, which may affect reproducibility and vesicle yield.
  - In this regard, we recommend using samples from each hemibrain processed independently (n = 2) from 2 animals (n = 4, N = 2).
  - Three indifferent preparations of antibody functionalization would then result in (n = 12, N = 3), providing sufficient biological and technical replicates to account for inter-sample variability and to perform robust statistical analysis.
  - This approach increases the reliability of observed SV recruitment patterns while minimizing bias from any single preparation or antibody coupling reaction.
  - Additionally, it is important to always image larger areas or acquire images spanning over a wider region to make sure that you are not skewing your data set while being biased during image acquisition.
- This protocol may not reliably detect low-affinity or extremely transient interactions, as weakly bound vesicles can be lost during the multiple washing steps.
- Lastly, while optimized for synaptic vesicles, adapting the method to other vesicle systems (e.g., exosomes, endosomes) may require re-optimization of labeling, antibody selection, and surface chemistry to maintain specificity and structural integrity.

## Troubleshooting 1

### Problem 1

#### Surface functionalization variability affecting capture protein immobilization

Variability in surface coating can lead to poor or uneven protein attachment, reducing vesicle recruitment efficiency. A well-coated coverslip should show a consistent low background puncta-like distribution, without patchy zones (Fig. 3, S3 and S2). This may result from suboptimal reagent freshness, improper pH during coupling reactions, antibody lot inconsistency, or coverslip contamination (please see SI, Fig. S2, S4). These inconsistencies can manifest as low or patchy fluorescent signals during imaging (related to preparatory step 1)

### Potential solution

- Always prepare fresh silane and crosslinker solutions immediately before use, especially for thiol-reactive reagents which degrade quickly in aqueous or oxygen-rich environments. Discard solutions after use; do not store overnight.
- Strictly control the pH (typically > 7.4 for maleimide coupling to NH groups) and temperature (RT) during surface functionalization steps. Buffering agents should be freshly prepared and acetone should be degassed (if necessary).
- Handle coverslips with powder-free gloves and avoid touching optical surfaces; clean coverslips using ethanol or opt for other protocols such as acid washing or plasma cleaning, if required. Use only high-quality microscopy-grade glass.
- Maintain consistency by using the same antibody lot for replicate experiments when possible. For new lots, validate binding capacity with control antigens, ELISA or vesicle-independent detection (e.g., fluorescent antibody binding assays).
- Include all validation control steps for background correction before vesicle incubation and compare. If background arises after SV incubation, then see troubleshooting problem 3.
- Make sure your protein of interest is indeed present in your sample and can be detected with the antibody functionalized on the glass surface, please run a WB or ELISA to examine this.

## Troubleshooting 2

### Problem 2

#### Low fluorescence signal after synaptic vesicle (SV) labeling

Low signal intensity can result from suboptimal dye concentration, incomplete or inefficient dye incorporation into vesicle membranes, degraded dye stocks, or compromised vesicle integrity during labeling or centrifugation. This hampers downstream imaging and quantification of vesicle recruitment (related to major step 2, section 3).

### Potential solution

- SynaptoRed is recommended for most systems due to high brightness and membrane specificity. However, BODIPY and Rhodamine-based dyes have also been used successfully. Choose dyes that match your microscope’s laser lines and emission filters. Test several options during initial optimization.
- Optimize exposure time and laser power settings as well as minimize dwell time on the sample during scanning.
- Use freshly thawed SV aliquots. Avoid multiple freeze-thaw cycles as they compromise membrane integrity, reducing dye uptake (Please see Fig. 2C-D).
- Use freshly prepared dye stocks, properly diluted to 1:1000 in the labeling buffer. Avoid using old or light-exposed dye stocks as degradation reduces efficiency.
- You can increase incubation time to enhance dye integration and optimization should be done quantitatively, please see, Fig. S4.

## Troubleshooting 3

### Problem 3

#### High background fluorescence, vesicle aggregation, or loss of vesicle integrity during incubation

Multiple variables during synaptic vesicle (SV) preparation and surface incubation can lead to poor imaging quality and compromised vesicle behavior. Elevated background fluorescence may result from non-specific binding, autofluorescence, or unbound dye due to insufficient washing (Fig. S4). Vesicle aggregation or bursting can occur from excessive SV concentration, prolonged incubation times, or mechanical and thermal stress during handling. Additionally, loss of vesicle integrity following trypsin treatment may arise from over-digestion of membrane proteins critical for vesicle stability. These issues commonly result in patchy, diffuse, or weak fluorescence signals that obscure accurate quantification of SV recruitment (related to major step 2, section 3, Fig. S4, S5).

### Potential solution

- Use optimal dilution of SR-loaded SV material since extremely high concentrations of material can cause vesicle crowding, membrane stress, and possibly non-specific binding.
- Handle SVs gently throughout preparation. Thaw on ice, avoid vertexing, and pipette slowly along the coverslip edge. Avoid forming bubbles. Limit incubation to 30–45 minutes at room temperature (∼22–25°C). Prolonged exposure increases the risk of vesicle rupture or surface crowding.
- Check on the efficiency of your washing steps using imaging buffer (e.g., flow buffer for 1 min, pause for 30 sec min, repeat) to remove unbound vesicles while preserving specific interactions.
- Acquire 6–10 background images from each coverslip prior to SV addition to allow for background correction and identification of surface irregularities. If the background is high before adding SVs, please stop imaging and start with preparing new coverslips again.
- Calibrate surface antibody density to ∼20–40 molecules/μm². Higher densities can lead to clustering *via* avidity effects, while extremely lower densities may reduce binding efficiency.
- For proteolytic treatments, optimize trypsin concentration (e.g., 5 µg/ml) and incubation time (e.g., 10 min). Immediately quench digestion with PMSF (1 mM) and 2% normal horse serum.
- Confirm vesicle integrity post-treatment via pellet size, protein quantification, or fluorescence intensity to ensure consistency across preparations. If needed, test alternative proteases with more specific activity.

## Troubleshooting 4

### Problem 4

#### Inconsistent control experiments by unmasking prey epitopes: Variability in blocking efficiency can obscure specificity assessment of vesicle recruitment

Inconsistent or incomplete blocking of synaptic vesicle (SV) surface proteins can arise from suboptimal antibody-to-vesicle protein ratios, poor incubation conditions, or limited epitope accessibility due to vesicle topology or protein conformation. These issues may lead to ambiguous interpretations of specificity in protein–protein interactions, making it difficult to distinguish true recruitment from background binding (related to major step 2, section 4 and 5).

### Potential solution

- Always prepare fresh antibody dilutions immediately before each experiment using sterile, filtered imaging buffer. Avoid repeated freeze-thaw cycles and store antibodies at recommended temperatures.
- Validate antibody activity and specificity before use *via* dot blots, immunocytochemistry, or vesicle pull-downs. Ensure the antibody recognizes the native conformation of the prey protein on SVs.
- Perform a dilution series (e.g., 1:1, 1:2, 1:3 relative to the standard working concentration) to determine the optimal antibody concentration. This can help identify both saturation thresholds and partial blocking regimes, which are useful for probing cooperative binding mechanisms.
- Include matched isotype control antibodies in parallel with each blocking experiment. These controls help distinguish specific inhibition of vesicle recruitment from background effects due to antibody concentration, Fc binding, or charge interactions.
- Ensure sufficient incubation time (15–30 min at room temperature with gentle mixing) to allow effective antibody binding to SV prey proteins. Avoid high temperatures or prolonged incubation that could destabilize vesicles.

## Troubleshooting 5

### Problem 5

#### Low fluorescence intensity or weak secondary antibody binding

Surface-exposed epitopes on recruited synaptic vesicles may be partially occluded or conformationally masked, resulting in inefficient secondary antibody access and reduced fluorescence signal (related to major step 3)

### Potential solution

- Use a biotin-conjugated primary antibody specific to one of the interacting proteins (e.g., SV-associated or prey protein). This circumvents epitope accessibility issues by targeting the antibody itself rather than relying on direct recognition of the native protein conformation on intact vesicles.
- Subsequently, you can label using fluorophore-conjugated streptavidin following biotinylated primary antibody incubation. This strategy benefits from the high affinity and small size of the streptavidin-biotin interaction, potentially improving signal intensity in sterically hindered environments.
- We avoid fixation, as we observed the loss of SR fluorescence (SI, Fig. S5). Keeping vesicles unfixed maintains their native state. Optionally, you could fix your coverslips using 4% PFA and then label the proteins of your interest. In our hands this resulted in a lot of background in the SR channel which one can choose not to image during data acquisition.

## Resource availability

- ***Lead contact****: Dr. Anne-Sophie Hafner*
- ***Technical contact****: Dr. Akshay Kapadia*
- ***Materials availability***: This study did not generate new, or unique reagents. All reagents are commercially available.
- ***Data and code availability***: All datasets generated during this study are made available upon request to the authors.

## Acknowledgments

- European Research Council (ERC) under the European Union’s Horizon 2020 research and innovation program - ‘MemCode’, grant 101076961 (AH)
- The authors thank all our lab members for fruitful discussions and helpful feedback.
- The authors acknowledge the support from General Instrumentation - Microscopy Core facility at Faculty of Science, Radboud University, Nijmegen, Netherlands; and thank Dr. Jelle Postma for his assistance with the TIRF microscopy set-up.

## Author contributions

AK conceptualized the project, performed and analyzed experiments and wrote the original draft of the manuscript. ASH supervised AK, acquired funding, and reviewed and edited the manuscript.

## Declaration of interests

The authors declare no competing interests.

## Notes

### Competing Interest Statement

The authors have declared no competing interest.

