## Supplementary Material file for "Single-molecule analysis of synaptic protein complexes and vesicle recruitment"

This SI file contains:

1. Supplementary protocols:
    a. Validation of methods and reagents (LOD determination of antibodies and proteins using ELISA)
    b. Workflow for data analysis using ImageJ plugins
2. Supplementary images depicting primary data (for other tested antibodies/conditions using this experimental protocol) as well as control data and images (artefacts and non-ideal images)

#### 1. a. Validation of LOD values using ELISA

- **Indirect ELISA:** Antigen → wash → primary antibody → wash → secondary antibody → wash → start reaction → stop reaction → read plate.
- **Sandwich ELISA:** Capture primary antibody → wash → block → antigen → wash → detection primary antibody → wash → secondary antibody → wash → start reaction → stop reaction → read plate.
- **Sandwich ELISA (customized):** Capture primary antibody → wash → block → antigen (capture protein, SYN) → wash antigen (prey protein, SV) → wash→ detection primary antibody (prey) → wash → secondary antibody → wash → start reaction → stop reaction → read plate.

- Primary antibody (capture/detection) incubation: Add 75 µl primary antibody solution (respective dilution in blocking buffer, 0.05 – 0.1 mg/well). for 2 h at RT or 16 h at 4°C, follow subsequent washing steps.
- Antigen incubation: Add 50-60 µl primary antibody solution (respective dilution in blocking buffer, 0.05 – 0.1 mg/well, optimized post pilot experiments). for 2 h at RT or 16 h at 4°C, follow subsequent washing steps.
- Washing steps: Remove residual liquid from the plate by gently tapping the plate (over tissue papers). Add 100 µl of 1x PBS and discard by tapping (2x quick washes and 2x washes with 5 min incubation intervals).
- Blocking step: Add 100 µl of blocking buffer (1 mg/ml BSA) per well and incubate for 2 h at RT. Tap the wells dry; washing steps are usually not required post blocking.
- Secondary antibody incubation: Add 50-60 µl streptavidin-conjugated-HRP or other HRP-conjugated antibodies (1:1000-5000) and incubate at RT for 2 h. Wash the wells thoroughly and tap dry; ensure there is no residual liquid remaining in the wells, since this would result in false positives.
- Reaction (start): Incubate 30 µl of 3,3′,5,5′-tetramethylbenzidine substrate (TMB; ThermoFischer Scientific, #34029) in each well at RT until sufficient blue colour developed (time ranged from 2-15 min depending on the different capture antibodies used in the experiment).
- Reaction (stop): Add 30 µl of stop solution (4 M $H_2SO_4$) to each well and measure absorbance values from each well on a plate reader at a wavelength of 450 nm and a background measurement at 620 nm. We used a Tecan multiwell plate reader.
- Analyze each sample in technical duplicate/triplicate wells in each experiment, as this would give you an idea of variation between wells as well as technical handling. If you see that for the same samples analyzed in duplicates/triplicates, values differ > 5%, please repeat the experiment.
- In certain cases, when you have a very low concentration of protein or an expensive antibody, an ELAST ELISA Amplification System (Revvity, #NEP116001EA) can be used as per the manufacturer's protocol to augment the signals *via* tyramide signal amplification.

#### 1. b. Particle analysis workflow for counting the number of recruited synaptic vesicles / FOV:

- Convert to 8-bit (Image → Type → 8-bit), as the system usually requires this.
- Adjust threshold (Image → Adjust → Threshold) to separate particles from background.
- Choose Default, Otsu, or Manual depending on your image or analysis parameters.
- Make sure only the round/circular particles of interest are red-highlighted.
- Go to Analyze → Analyze Particles…. The important settings:

  Size (µm² or pixels) -Sets the minimum and maximum particle area to include.

- Since we are analyzing round particles of the SynaptoRed labelled synaptic vesicles, please set,
  - Minimum: ~0.02–0.05 µm² (exclude noise signal/ extremely small particles)
  - Maximum: depends on avoiding aggregates (maybe <0.5–1 µm²).
- The values depend on your calibration and quantification p[parameters (set under Analyze → Set Scale).
- Additionally, you can also set for circularity (0.00–1.00), wherein 0.00 = any shape and 1.00 = perfect circle.
  - For synaptic vesicles, you might use 0.6–1.0 to target near-spherical particles and avoid irregular debris.
- Outlines – draws contours of detected particles.
- Masks – creates a binary mask of detected particles.
- Nothing – just outputs the results table.
- Checkboxes (make sure to select based on the desired analysis output)
  - Display Results – outputs particle measurements (area, mean intensity, etc.).
  - Summarize – gives total count, total area, and mean size.
  - Exclude on Edges – avoids counting particles touching image borders.
  - Add to Manager – stores selections for later review.
- Inspect detected particles visually to ensure threshold and selection criteria match actual vesicles.
- Depending on the images you acquire, sometimes aggregates may be counted, adjust the max size or circularity, so as to avoid detection of false positives.
- For consistency, apply the same settings to all images in the particular dataset of the same experiment (use Process → Batch). This would strictly apply conditions with and without blocking/proteolytic cleavage, etc.

### Supplementary images

**Figure legends:**

**Figure S1. Another example of antibody thiolation and validation of capture efficiency.**

(A) Validation of RIM1 antibody reactivity post-thiolation. (i) SDS-PAGE under various reducing and non-reducing conditions showing heavy and light chains detected by Coomassie, Ponceau, and western blot (WB) with anti-rat secondary. N = 2. (ii) Quantification of antibody reactivity by WB and modified ELISA reveals that SATA thiolation (condition 6) did not significantly impair antibody function, and reactivity is preserved after deprotection (condition 7). WB, N = 2, ELISA, N = 3.

(B) RIM1 antibody reactivity in indirect and sandwich ELISA assays. (i–ii) RIM1 antibodies bind to RIM1 protein in synaptosome lysates in a dose-dependent manner, with or without thiolation. Examined using indirect ELISA. (iii) Sandwich ELISA showing capture of Rab3a to RIM1 protein immobilized *via* antibody without (iii; n = 4, N = 2) and with thiolation steps (iv; n = 6, N = 3). Here, the difference in binding and protein capture efficiency indicates RIM1 antibody activity is slightly altered post-modification.

(C) Western immunoblotting images depicting antibody modification (both RIM1 and STX1A) using partial thiolation methods - βME treatment for different time points) completely abrogated antibody reactivity.

**Figure S2. Optimization of surface functionalization with an antibody for SIM-Pull assays.**

(A) Schematic of surface activation and functionalization strategy. Coverslips are sequentially modified by aminosilanization (APTES), maleimide activation (Sulfo-SMCC), and covalent coupling of thiol-modified antibodies *via* SATA and $NH_2OH$. The final antibody-conjugated surface is used for vesicle or protein pulldown in the SIM-Pull assay.

(B) TIRF images show the effect of antibody coating concentration and agitation on spot density and uniformity. Surfaces were incubated with increasing concentrations (1, 2 and 5 µg/mL) of antibody without (middle row) or with shaking (bottom row). Activation of the glass surface is necessary for effective functionalization. Insets show higher-magnification views (scale bars: large panels = 10 µm; insets = 0.5 µm). n = 2.

(C) Antibody specificity screen using five different antibodies (AB#1–AB#5) at a fixed concentration (0.5 µg/mL). AB#1–AB#4 show effective surface binding and discrete spot formation. AB#5 yields a low signal, and the no-antibody control shows negligible background. Scale bars: 10 µm. n = 2.

(D) Multichannel TIRF imaging of an ideal AB#1 functionalized surface. Channel 1 (Ch1, 488 channel) shows a robust spot signal from the capture antibody, while Channels 2–4 (Ch2–Ch4, 405, 555, 647 channels, respectively) show minimal or no signal, indicating specificity and minimal crosstalk across channels. Scale bars: left panel = 10 µm; right panels = 0.1 or 0.5 µm. Images are representative of multiple experimental replicates performed during the entirety of this manuscript.

**Figure S3. Quantification of particle density and variability across experimental conditions post surface functionalization.**

(A-C) Representative thresholded images (STX1A, STX1, and Syb; data depicted in Fig. 3A-C) used for analyzing particles of immunoisolated proteins labelled with fluorescent antibodies: (A) STX1A only (*red*), (B) STX1A (*red*) and STX1 (*green*), and (C) STX1A (*red*) and Syb (*blue*). Numbers in images indicate percentage of area covered with particles per FOV (mean ± SD). Right, quantification of the number of detected particles per field of view (FOV) from independent experimental replicates. Data are shown as mean ± SD; individual points represent independent measurements from separate FOVs. Scale bars, 5 µm. n = 6, N = 4 (indicated as rep. 1a-b and 2a-b, respectively)

**Figure S4. Immunocytochemistry to probe capture-prey (STX1A-Syb) protein complexes tethering SR-SV**

(A) Representative overview images post-ICC showing SR-SV (*magenta*) and STX1A (*red*), and Syb (*green*) channels. Zoomed-in images are depicted in the figure. 5B. N = 3.

(B) Representative overview images post-ICC showing SR-SV (*magenta*) and RIM1 (*red*), and Rab3a (*green*) channels. N = 3.

(C) Zoomed-in panels depicting the tethering of SV show that it is indeed *via* the interaction of RIM1 and Rab3a. The data is representative of three independent experiments.

**Figure S5. Representative examples of ideal and non-ideal SV TIRF images**

Representative images showing synaptic vesicle (SV) preparations labelled with SynaptoRed (SR-SV, *magenta*). Top row: During SV incubation, ideal preparations display well-dispersed SVs (*left*), whereas non-ideal preparations exhibit excessive concentrations and aggregated SVs (*right*). Bottom row: After washing steps, ideal conditions yield well-separated, single tethered SVs (*left*), while non-ideal conditions result in aggregated SVs with larger diameters, which could also result from sterically bound SVs remaining due to reduced washing steps (*right*); scale bars, 5 µm. Please note, all images depicted here are at saturated fluorescence intensities for better visualization.

**Figure S6. Representative examples of ideal and non-ideal SV TIRF images post-ICC and washing steps**

(A) Representative single-molecule TIRF images of SR-SV (*magenta*) showing effects of common preparation errors. Excessive washing can result in SV loss (*left*) or loss of SR fluorescence (*middle*), while the use of a non-isotonic buffer disrupts SV integrity (right), leading to increased background fluorescence. A decrease in fluorescence signals could also be a result of photobleaching the dye due to strong laser power. Scale bars, 5 µm.

(B) Examples of ideal and non-ideal images following ICC and washing. Top row: Ideal images display well-dispersed, intact SVs (*white*) with clear co-localization to capture protein x (*green*). Middle row: Non-ideal but usable images show SVs (SR fluorescence is slightly lost), but images are still suitable for protein localization or quantification (protein y, *blue*).

Bottom row: Non-ideal images exhibit improper labelling of protein z (*magenta*), necessitating optimization for ICC labelling of protein. Merged channels are shown in the leftmost panels; SV-only channels in white; protein-specific ICC channels in color. Scale bars, 10 µm (*leftmost panels*) and 0.5 µm (*zoomed panels*).

**Figure S1.**

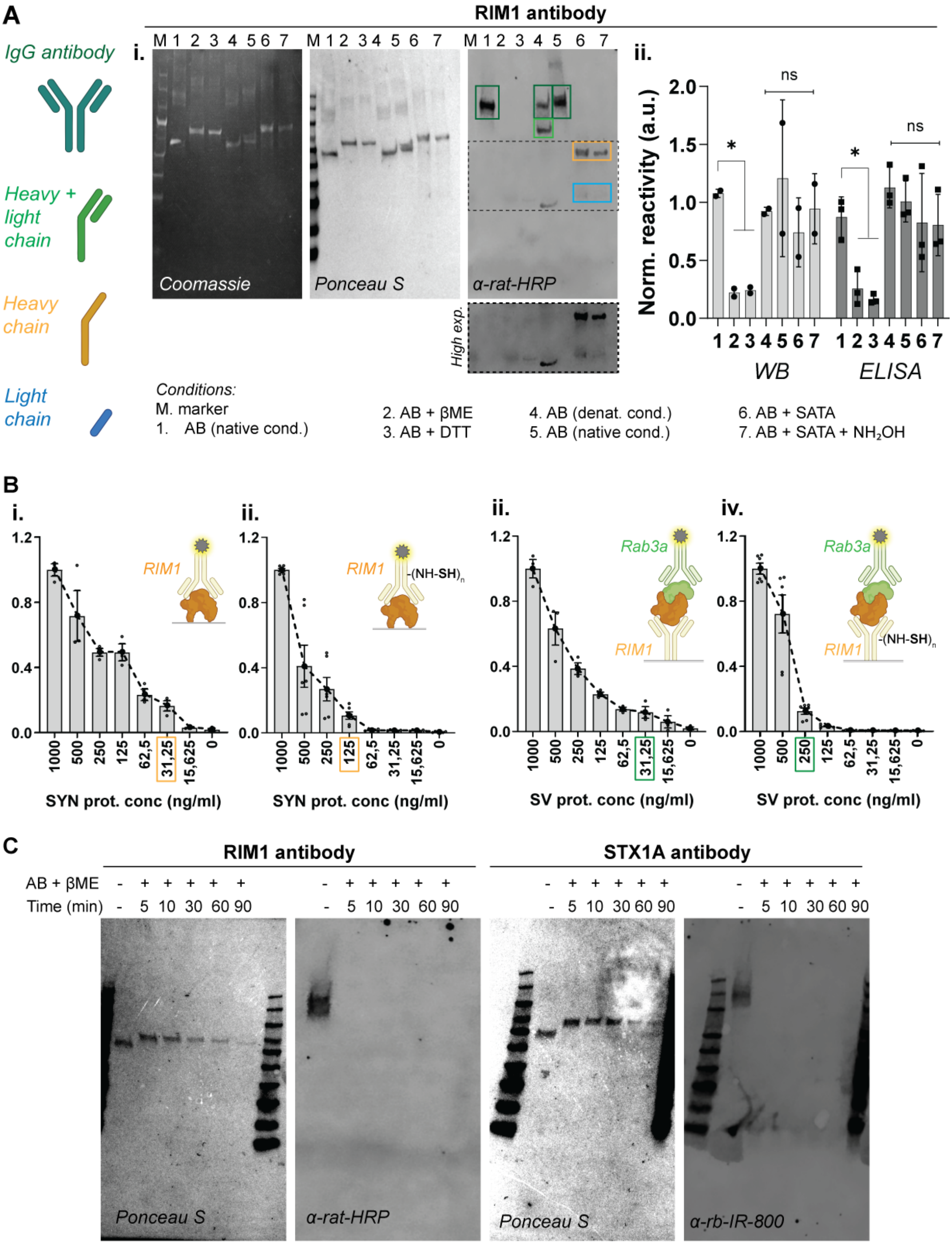
A
RIM1 antibody
IgG antibody
Heavy + light chain
Heavy chain
Light chain
i.
M 1 2 3 4 5 6 7
Coomassie
Ponceau S
α-rat-HRP
High exp.
ii.
Norm. reactivity (a.u.)
2.0
1.5
1.0
0.5
0.0
ns
*
1 2 3 4 5 6 7
WB
ELISA
Conditions:
M. marker
1. AB (native cond.)
2. AB + βME
3. AB + DTT
4. AB (denat. cond.)
5. AB (native cond.)
6. AB + SATA
7. AB + SATA + NH₂OH
B
i.
ii.
ii.
iv.
RIM1
-(NH-SH)n
Rab3a
1.2
0.8
0.4
0.0
1000
500
250
125
62,5
31,25
15,625
0
SYN prot. conc (ng/ml)
SV prot. conc (ng/ml)
C
RIM1 antibody
STX1A antibody
AB + βME - + + + + +
Time (min) - 5 10 30 60 90
Ponceau S
α-rat-HRP
α-rb-IR-800

**Figure S2.**

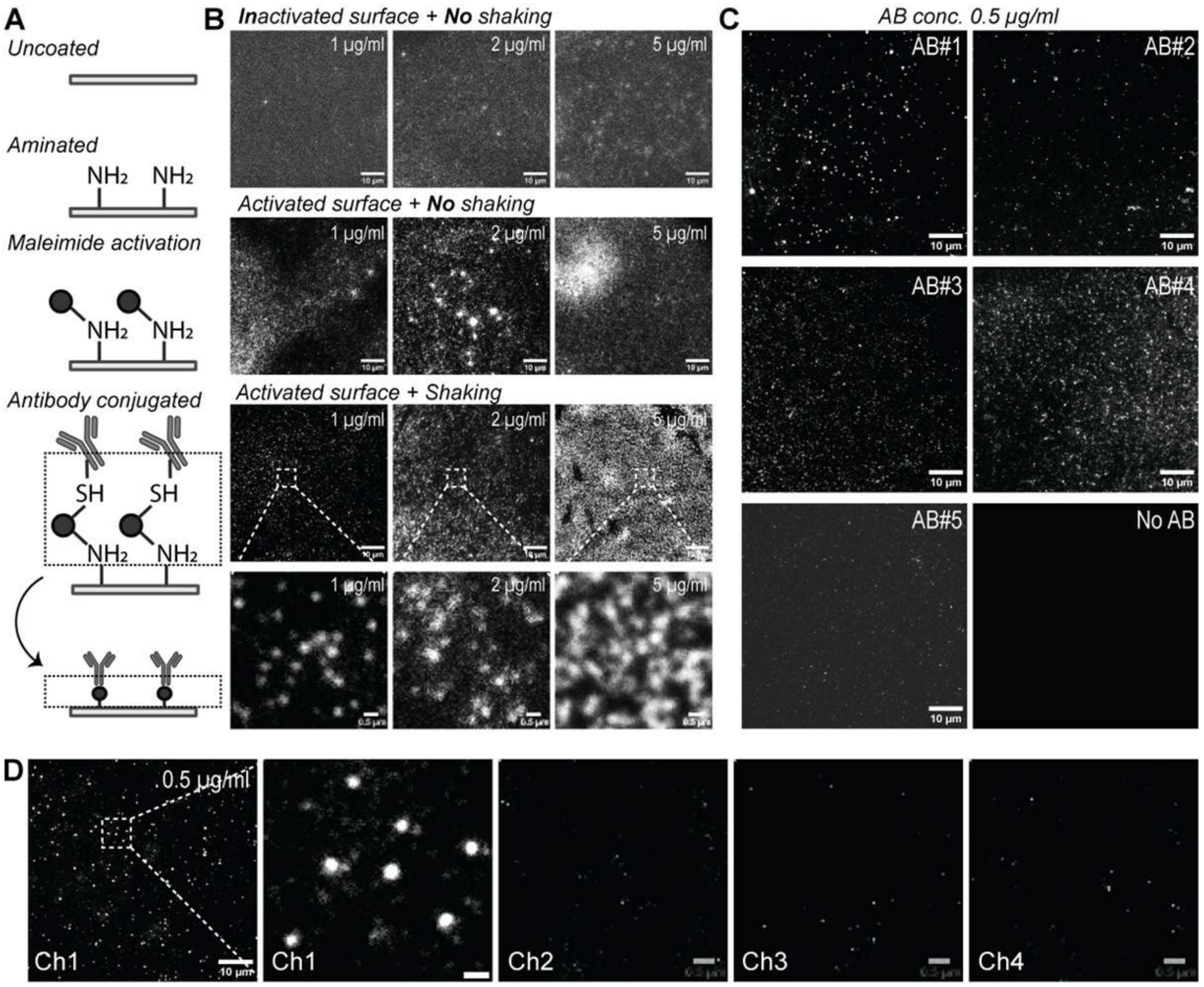
A
Uncoated
Aminated
NH2 NH2
Maleimide activation
Antibody conjugated
SH SH
B
Inactivated surface + No shaking
1 μg/ml
2 μg/ml
5 μg/ml
Activated surface + No shaking
Activated surface + Shaking
10 μm
0.5 μm
C
AB conc. 0.5 μg/ml
AB#1
AB#2
AB#3
AB#4
AB#5
No AB
D
0.5 μg/ml
Ch1
Ch2
Ch3
Ch4

**Figure S3.**

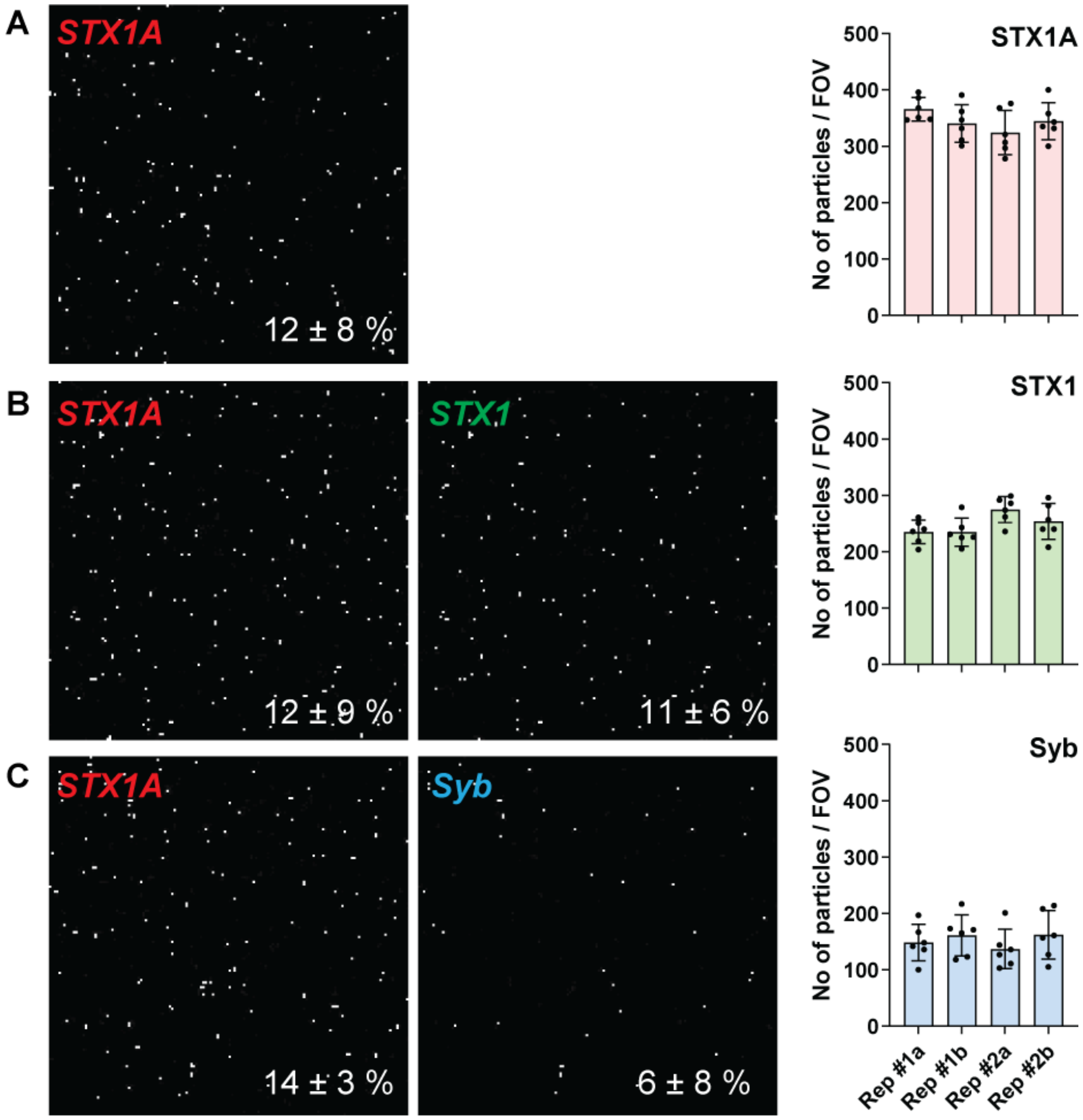

A
STX1A
12 ± 8 %
B
STX1A
12 ± 9 %
STX1
11 ± 6 %
C
STX1A
14 ± 3 %
Syb
6 ± 8 %
STX1A
STX1
Syb
No of particles / FOV
500
400
300
200
100
0
Rep #1a
Rep #1b
Rep #2a
Rep #2b

**Figure S4.**

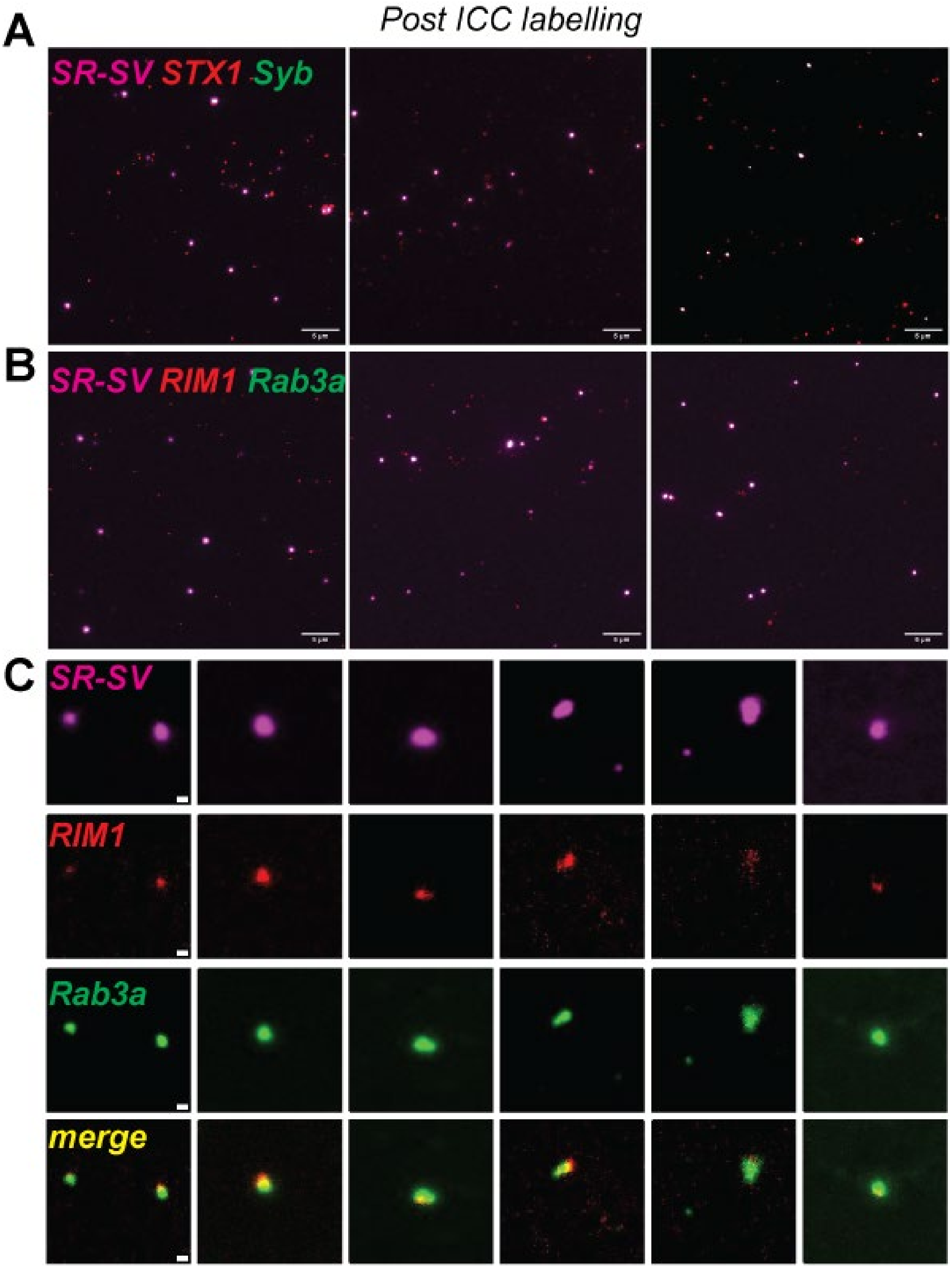

Post ICC labelling
A
SR-SV STX1 Syb
B
SR-SV RIM1 Rab3a
C
SR-SV
RIM1
Rab3a
merge

**Figure S5**

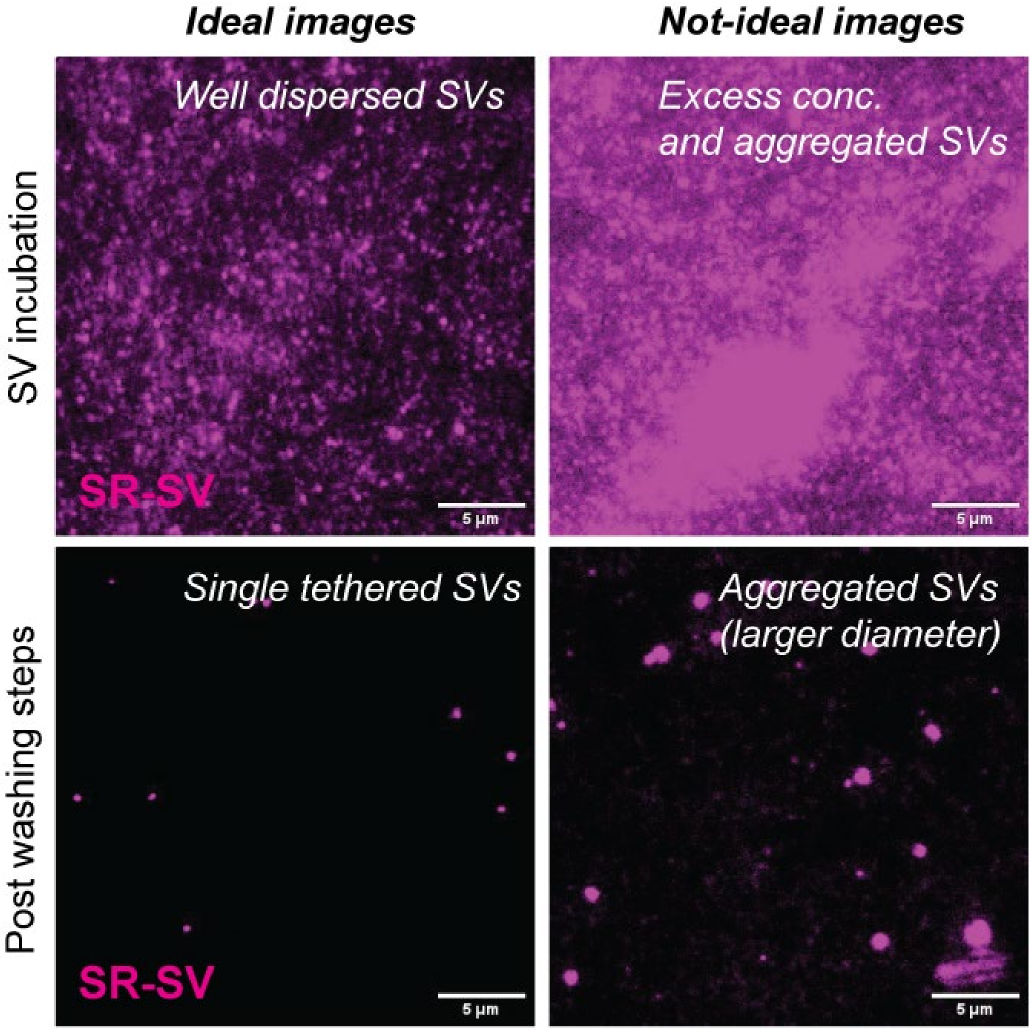
Ideal images
Not-ideal images
SV incubation
Post washing steps
Well dispersed SVs
Excess conc.
and aggregated SVs
Single tethered SVs
Aggregated SVs
(larger diameter)
SR-SV
SR-SV
5 µm
5 µm
5 µm
5 µm

**Figure S6.**

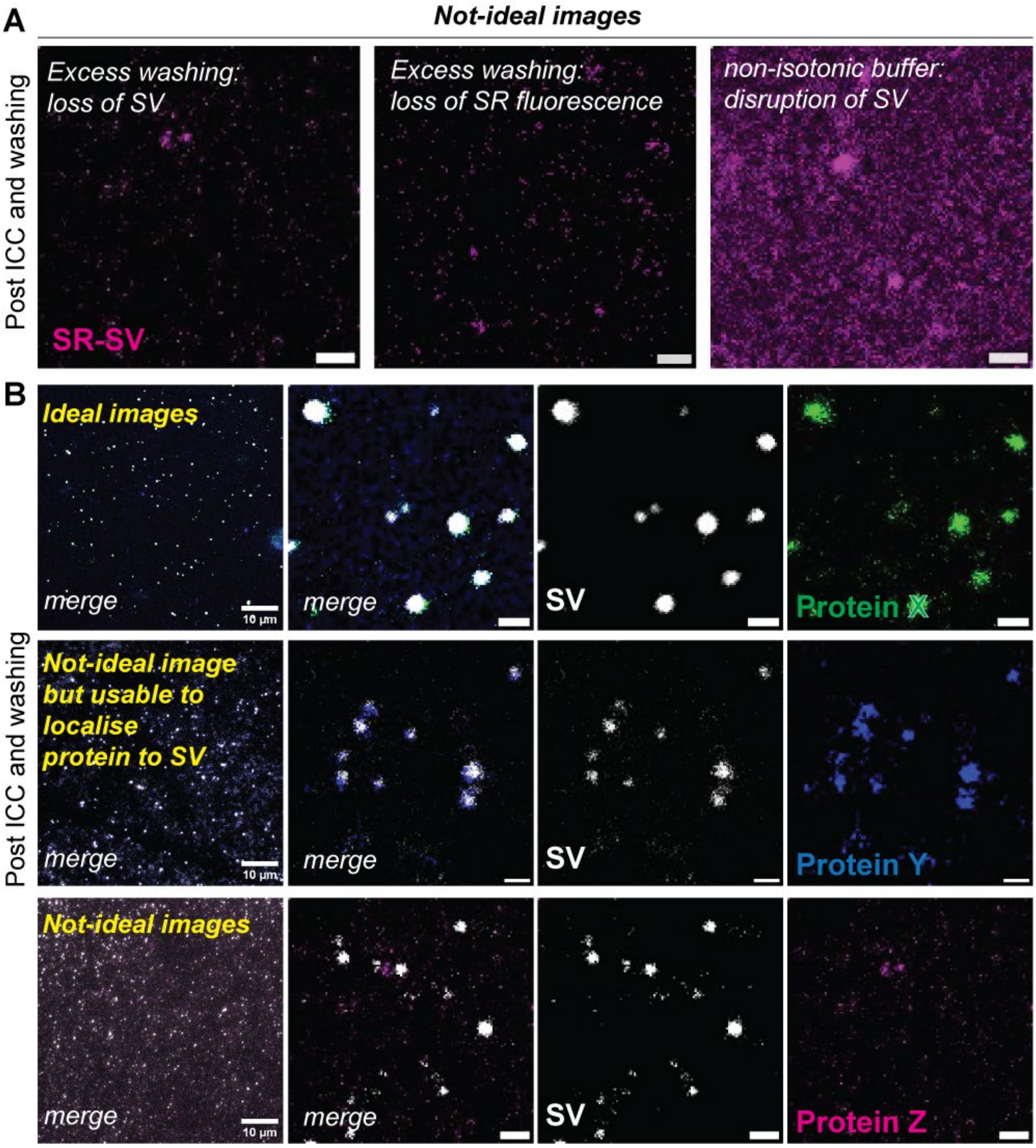
A
Not-ideal images
Post ICC and washing
Excess washing: loss of SV
SR-SV
Excess washing: loss of SR fluorescence
non-isotonic buffer: disruption of SV
B
Ideal images
merge
10 µm
merge
SV
Protein X
Post ICC and washing
Not-ideal image but usable to localise protein to SV
merge
10 µm
merge
SV
Protein Y
Not-ideal images
merge
10 µm
merge
SV
Protein Z
